# Fast Optimization of Robust Transcriptomics Embeddings using Probabilistic Inference Autoencoder Networks for multi-Omics

**DOI:** 10.1101/2025.11.16.686778

**Authors:** Ning Wang, David Turner, Hannah Feinberg, Victor Eduardo Nieto Caballero, Dan Yuan, Nathaniel Scott, Christopher Cardenas, Michael DeBerardine, Shu Dan, Lakme Caceres, Jessica Schembri, Zizhen Yao, Changkyu Lee, Jonathan W. Pillow, Fenna M. Krienen

**Affiliations:** Princeton Neuroscience Institute (PNI); Princeton Program in Quantitative and Computational Biology (QCB); Allen Institute for Brain Science (AIBS); Princeton Center for Statistics and Machine Learning (CSML); Princeton Program in Applied and Computational Mathematics (PACM)

**Keywords:** Single cell RNA sequencing, Multi-omics, Integration, Cross-species atlases, Transcriptomics, Spatial Transcriptomics, Deep Learning, Methods Development

## Abstract

Advances in single-cell genomics technologies enable the routine acquisition of atlases with millions of cells. These datasets often include multiple covariates, such as donors, sequencing platforms, developmental timepoints, and species, which provide new opportunities for discovery and new challenges. To mitigate unwanted sources of variation, dataset integration is the starting point for most analyses. However, existing methods struggle with integrating large complex datasets. To address these limitations, we developed PIANO, a variational autoencoder framework that uses a negative binomial generalized linear model for stronger batch correction, and code compilation for ten times faster training than existing tools. We first demonstrate performant integration compared to commonly used methods on single-species datasets. We then show PIANO enables superior analyses of multiple atlases, solving challenging integration tasks across sequencing platforms, development, and species, while simultaneously preserving desired biological signals. Our contributions include a novel, high-performance integration method and recommendations for integration applications.

## INTRODUCTION

Single cell sequencing enables the creation of cross-species and cross-modality atlases with millions of cells and nuclei, such as whole mouse and human brain atlases^1,2^. This powers large consortium-based efforts to map the cellular composition of entire organs or even organisms, such as those from the BRAIN Initiative Cell Atlas Network (BICAN), Human Cell Atlas (HCA), and Chan Zuckerberg Initiative (CZI). Comparing data from these large studies enables exploration of the evolutionary conservation and divergence of cell types across multiple species, developmental timepoints, and modalities^1–4^. Consequently, such atlases now scale to tens of millions of cells, which calls for powerful new tools for analyzing the data.

While combining many datasets enables novel biological insights, single cell sequencing datasets are still heavily impacted by data sparsity and multiple sources of variation. Sources of variation such as the tissue donor and sequencing platform – often referred to collectively as batch effects – confound biological discoveries and make it difficult to compare cell types across experimental conditions. Therefore, one goal of data integration is to find a more informative, low-dimensional representation of the original data. Ideally, the integrated latent space representation should remove batch effects while preserving relevant biological information (e.g. cell types).

To enable robust comparisons across datasets, multiple methods have been developed for projecting single cell or single nucleus sequencing data to a shared latent space embedding^5^. One commonly used approach is Seurat’s canonical correlation analysis (CCA), which finds anchors between batches by identifying pairs of similar cells across datasets^6^. However, using pairwise anchors may over-mix data in situations with imbalanced cell type distributions between pairs. While Seurat may be sufficient for simpler tasks, pairwise anchor calculations scale quadratically with the number of batches^6,7^. Moreover, the sparse matrices used in Seurat’s implementation cannot handle more than 2^31^ - 1 non-zero gene counts in memory, which further limits its scalability. Another popular method, Harmony, uses expectation-maximization (EM) to perform batch correction^5^. However, recent benchmarking studies suggest Harmony also struggles on complex integration tasks^7^.

An alternative solution is to use a variational autoencoder (VAE)^8^ for integration. A VAE is a powerful, non-linear, generative model that uses an encoder-decoder architecture to explain high-dimensional data using a low-dimensional latent variable. Through dimensionality reduction, the VAE extracts the most meaningful information from the data, such as information about cell type clusters. The generative component of the model, or decoder, describes a transformation from this low-dimensional latent space to the high-dimensional space of the raw counts data. The other component of a VAE is the recognition model, or encoder, which enables inference of the latent variables from an observed single cell dataset. When used for integration, the VAE trains on data from multiple batches and integrates the data by performing batch correction, minimizing the differences between batches.

One popular VAE method, scVI, integrates single cell datasets using a negative binomial (NB) distribution to account for over-dispersion in the raw gene counts^9^; scVI performs batch correction by conditioning the over-dispersion term for each gene on the primary source of variation^9^. Although the scVI decoder can take additional covariates as input, each NB distribution is only conditioned on one primary batch key. Moreover, despite benefits from graphics processing unit (GPU) acceleration as an end-to-end trained neural network, scVI’s training time scales linearly with the size of the data^7,9^. As a result, scVI may take days to train on large datasets with millions of cells, which delays downstream analyses.

To address these limitations, we introduce PIANO: <u>P</u>robabilistic Inference <u>A</u>utoencoder <u>N</u>etworks for multi-<u>O</u>mics, a novel variational autoencoder framework for inferring integrated latent space representations for single cell transcriptomics data. We utilize a generalized linear model (GLM) to fit a negative binomial (NB) distribution for each gene to address unwanted sources of variation. Rather than relying on a primary batch key like current VAE methods^9^, PIANO accounts for all covariates using its NB-GLM. PIANO integration achieves superior or comparable biological conservation, which measures preservation of cell type clusters, with stronger batch correction, which measures mixing of data across batches, on multiple integration tasks, as assessed by commonly used integration benchmarks. Moreover, our light-weight code implementation that utilizes code compilation leads to substantially faster training speed, resulting in up to 10 times faster training than current VAE models.

## RESULTS

### NB-GLM variational autoencoder enables integration of gene count data

To enable integration by reducing batch effects, PIANO learns a latent representation of the data using a variational autoencoder (VAE)^8^ that incorporates external covariates (Figure 1). PIANO uses a deep neural network (DNN) inference model, or "encoder", to perform a non-linear mapping from vectors of gene-count observations to lower-dimensional latent variables, learning an approximate distribution over the latent representation for each cell (Figure 1). To describe gene count data, PIANO uses a generative model, or “decoder”, which uses another DNN to perform a non-linear mapping between latent variables and their covariates to the parameters of a negative binomial (NB) distribution that models the original observations. Due to the information bottleneck imposed by low-dimensionality, the latent variable must learn the most informative representation of the data to help the decoder model the original data. The final layer of the decoder incorporates a NB-GLM — a generalized linear model (GLM) with NB observations for each gene, which combines its DNN outputs with linear weights on the library size (i.e., total counts per cell) and batch-related covariates, such as donor or species (see Methods, Supplemental Table 1). Since these covariates are explicitly provided to both the decoder and to the GLM, the latent variable itself does not need to inform the decoder about covariates for modeling the data and instead encodes cell-type information with greatly reduced batch effects. This GLM framework accounts for multiple overlapping sources of variation, which offers greater flexibility and stronger batch correction than scVI’s conditional batch key (see Methods).

**Figure 1:**
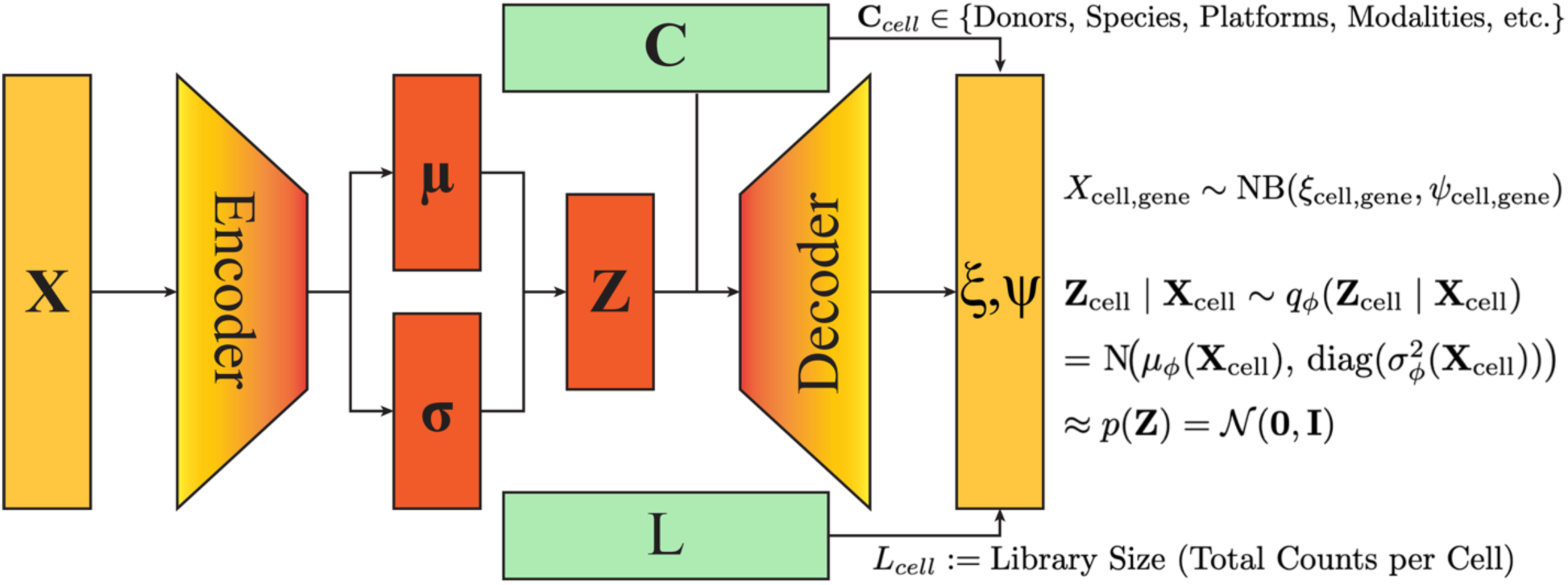
PIANO uses a variational autoencoder (VAE) architecture. The encoder finds a low-dimensional representation Z of the data X. The decoder uses Z, the covariates C, and the library size L to model the data. The encoder-decoder architecture enables unsupervised training of the VAE. The decoder uses C and L in a generalized linear model (GLM), which enables PIANO to perform batch correction for data integration.

### PIANO integrates across single-species donor and technical variability

We first demonstrated PIANO integration on five published single-species cases. Datasets varied in the number of cells (16k-373k), cell types (11-81) and tissues, but all contained known batch effects, such as donors and sequencing platforms^10–21^ (Figure 2). To quantify integration performance, we used two metrics for biological conservation: normalized mutual information (NMI) and adjusted Rand index (ARI), and two metrics for batch correction: K-nearest-neighbors batch effect test (KBET) and integration local inverse Simpson’s index (iLISI) (Methods). PIANO has a strong overall integration performance compared with other integration methods (Seurat, scVI, and Harmony) and an unintegrated baseline using principal component analysis (PCA) (Figure 2a, Supplemental Table 2). Due to stochasticity in model initialization and the relatively small dataset sizes, both the PIANO and scVI integrations were repeated five times, and the integration benchmarking metrics were averaged over the five integrations. For average biological conservation between NMI and ARI, PIANO had comparable performance to scVI and generally stronger performance to Seurat (Figure 2b, Supplemental Table 3). For average batch correction between KBET and iLISI, Seurat generally had the strongest performance, followed by PIANO (Figure 2c, Supplemental Table 4). However, Seurat may have overcorrected for batch effects, which may have contributed to Seurat’s underperformance in biological conservation (Figure 2b). While Seurat tended to overintegrate, Harmony and PCA had generally lower integration performance for most datasets across all metrics (Figure 2a-c). This can be appreciated from the uniform manifold approximation and projection (UMAP) visualizations of the respective latent spaces, where over-integrated data results in blended cell type clusters (Supplemental Figures 1-5). In contrast, Harmony tends to underintegrate the data (Supplemental Figures 1-5). Individual integration UMAPs and benchmarking scores can be found in supplemental figures and tables (Supplemental Figures 1-5, Supplemental Tables 5-9).

**Figure 2:**
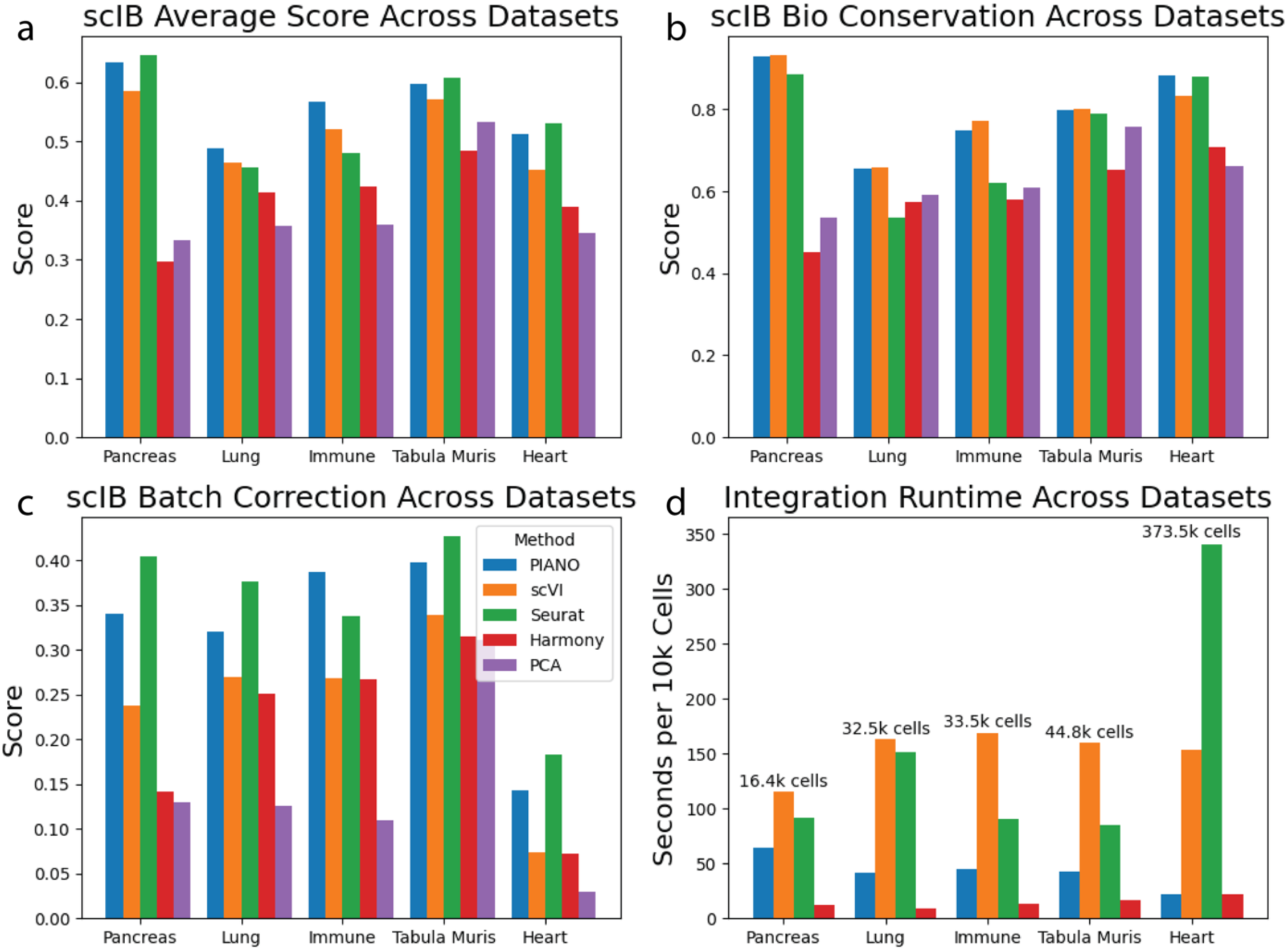
Benchmarking performance across methods. PIANO has superior or comparable average integration benchmarking performance (a), biological conservation (b), and batch correction (c) across datasets compared to currently available methods. PIANO has substantially lower runtime costs, including compilation time, compared to all methods except Harmony for all datasets (d).

For the smaller datasets, scVI and Seurat had similar runtimes, but on the larger heart dataset, Seurat took over twice as long to run as scVI (Figure 2d). In contrast, PIANO was faster to train than all integration methods besides Harmony for every dataset; PIANO was still faster even when including compilation time, which accounted for the majority of the total training time on the smaller datasets (Figure 2d). On the larger heart atlas (n=343.5k cells), where compilation time accounts for a smaller proportion of training time, PIANO using one worker in its GPU memory mode (Methods) was seven times faster than scVI using eleven workers and 15.5x faster than Seurat (Figure 2d); both PIANO and scVI models were trained using the same type of hardware: one NVIDIA A100 GPU with 40Gb of GPU memory. Runtimes are in Supplemental Table 10.

### PIANO integrates atlas-level data across the primate basal ganglia

To better understand the behavior of VAE models for single cell integration, we investigated multiple variations of feature selection and model design using a large multi-species brain cell atlas of the basal ganglia (Figure 3), acquired as part of the BRAIN Initiative Cell Atlas Network (BICAN) (Johansen et al., 2025). The Human and Mammalian Brain Cell Atlas (HMBA) contains RNA sequencing data from nearly 2 million single nuclei sampled across human, macaque, and marmoset basal ganglia structures. Each species’ data were originally computationally integrated using scVI to generate a consensus cell taxonomy at two hierarchical levels: cell Class (12 Classes) and cell Group (61 Groups) (Johansen et al., 2025).

**Figure 3:**
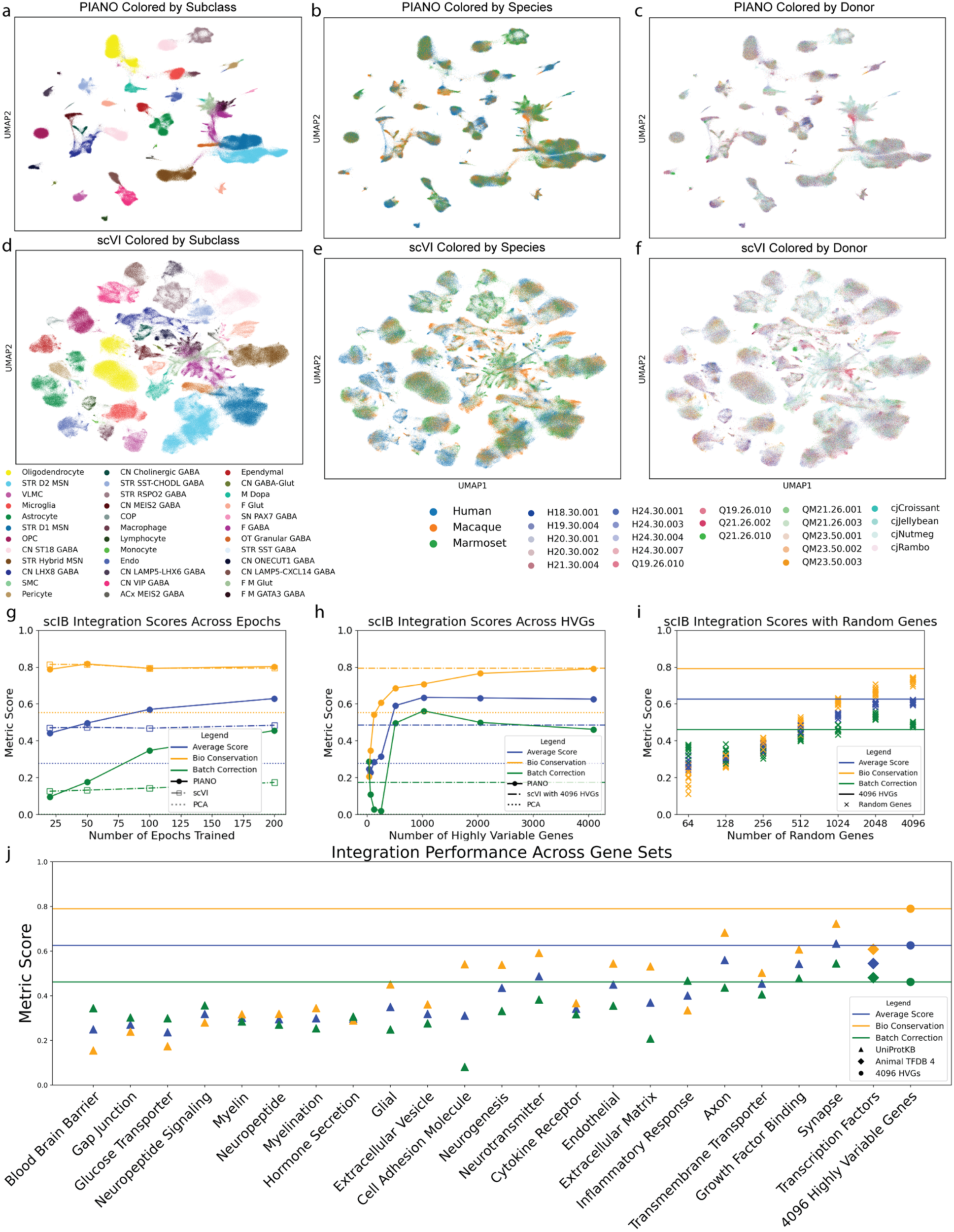
Cross-species integration of primate basal ganglia. PIANO enables robust integration of primate basal ganglia, colored by Subclass (a), species (b), and donor (c) compared to scVI, colored by Subclass (d), species (e), and donor (f). PIANO has comparable biological conservation as scVI and stronger batch correction over epochs (g). We find that using 4096 highly variable genes offers good integration performance (h) and better performance than random genes (i) or functional gene sets, such as transcription factors or neurotransmitter-related genes (j).

To balance the number of cells between species, we subset the data to include up to 6,400 cells per Group for each species. Using the Group labels to assess biological conservation, PIANO performed a better separation of cell types and mixing between donors and species (Figure 3a-c) compared to scVI (Figure 3 d-f). We found that a model architecture with 3 layers, 256 hidden nodes, and 32 latent dimensions had a well-balanced integration performance (Figure 3g, Supplemental Figure 6). As the number of training epochs increased, the batch correction improved while the biological conservation remained stable (Figure 3g). In contrast, scVI had a less noticeable improvement in integration performance over training (Figure 3g). While PIANO had similar biological conservation of Groups to scVI, PIANO had substantially higher batch correction across species (Figure 3g). Additionally, both PIANO and scVI outperformed using a PCA latent space, which offered no batch correction (Figure 3g).

Gene selection is an important parameter for integration tasks, particularly when integrating datasets across species. We first subset the data to 1:1 homologous genes^22^ and then selected the top highly variable genes (HVGs) across species for integration using the "Seurat V3" flavor for highly variable gene selection, using species as the "batch_key". We performed a hyperparameter sweep over the number of top HVGs and found that using 4096 HVGs for integrating the HMBA datasets had robust performance (Figure 3h). Surprisingly, the same number of randomly selected genes had comparable integration performance with similar batch correction and slightly worse biological conservation (Figure 3i). This suggests that PIANO can robustly integrate data given sufficient high-quality, gene expression training data.

Several atlas studies have prioritized functional gene sets to construct single-species cell type taxonomies. The Allen Institute whole mouse brain (WMB) atlas compared cross-validation accuracy of clustering using all cluster-wise differentially expressed genes (n=8,460 genes) against several functional gene sets including transcription factors (TFs, n=534 genes), cell adhesion molecules (n=857 genes) and other functional classes (n=541 genes)^2^. This analysis demonstrated that cross-validation accuracy using TFs was comparable to all expressed genes for discriminating between 338 subclasses of cells^2^. In the WMB taxonomy, the Subclass is considered an intermediate level of the atlas^2^ and approximates the Group level in the HMBA dataset (Johansen et al., 2025). We examined whether functional gene classes performed similarly to large sets of highly variable genes for cross-species integration (Figure 3j). The larger functional gene sets, such as the orthologous synapse related genes from the UniprotKB^23^ (838 genes) and orthologous transcription factors from AnimalTFDB4^24^ (1,119 genes) showed similar performance to the 4096 highly variable genes and outperformed the orthologous extracellular matrix genes (414 genes) (Figure 3j). This suggests that functional gene sets that are sufficiently large contain enough information to capture the nuances between cell type clusters (Figure 3d-g). UMAPs for PCA can be found in Supplemental Figure 7, and specific benchmarking scores can be found in Supplemental Tables Supplemental Table 11-13.

One mode of evolutionary divergence could come as complete gain or loss of a cell type^25^. As such, an ideal cross-species integration should preserve species-specific cell type populations^26^. We performed two *in silico* perturbation experiments on evolutionarily conserved cell types to determine whether or not PIANO and scVI would overcorrect for species effects. To determine if a species-specific cell type would be merged into a related cell type shared with other species, we first withheld only the hybrid medium spiny neurons (Hybrid MSNs) Subclass from the non-human primate datasets, such that only the human basal ganglia dataset retained them. Both PIANO and scVI kept these Hybrid MSNs separate from the related D1 and D2 MSNs (Supplemental Figure 8). We then withheld only the non-human primate D1 MSNs. Since the D1 MSNs are much more transcriptionally similar to the D2 MSNs than are the Hybrid MSNs, this constitutes a more difficult perturbation. Initially, both PIANO and scVI mixed the D1 and D2 MSNs, resulting in overlapping cell type clusters (Supplemental Figure 9); however, by lowering the value of the beta hyperparameter while training PIANO (see Methods), the D1 MSNs were re-separated from the D2 MSNs (Supplemental Figure 9). These results suggest PIANO may require tuning to better retain dataset-specific populations, depending on the relative similarity between shared and imbalanced cell populations across datasets.

### Integration methods are challenged by datasets with large noise effects

Cross-species integrations become more challenging with increased evolutionary divergence, particularly when each species’ dataset also contains major within-dataset covariates such as different developmental timepoints. To assess a more complex scenario we evaluated PIANO integration performance between the mouse cell atlas (MCA)^27^ and human cell landscape (HCL) datasets^28^ (Figure 4). Not only did these datasets have many cell types across species and developmental time points, they also had low signal-to-noise, with fewer counts captured per cell compared to more recently collected datasets^26^ due to using an older Microwell-Seq platform^27,28^. Consequently, we found it difficult to maintain separable cell type clusters while mixing across species for PIANO (Figure 4a-b) and scVI (Figure 4c-d, Figure 4 S1), which resulted in lower overall integration performance than previous integration examples. PIANO and scVI had similar biological conservation scores, and scVI performed very little batch correction (Figure 4e). Moreover, we found that PIANO trained up to ten times faster than scVI as the number of cells increased (Figure 4f). UMAPs for PCA (including the cell types legend) can be found in Supplemental Figure 10, and benchmarking metrics and runtimes can be found in supplementary tables (Supplemental Table 14-15).

**Figure 4:**
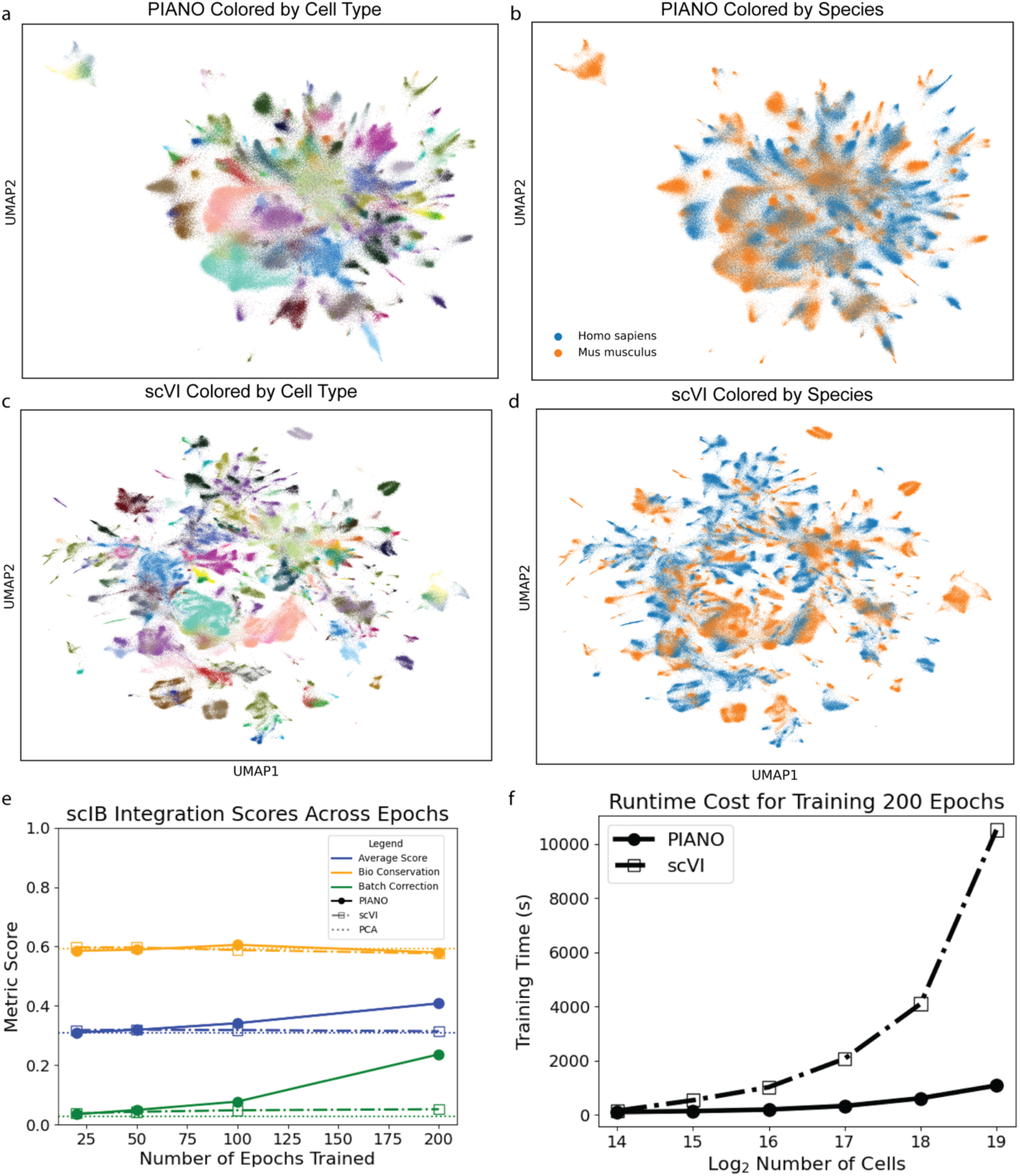
Mouse cell atlas and human cell landscape integration. All methods failed to integrate the mouse cell atlas and human cell landscape atlases, as seen in the PIANO integration uniform manifold approximation and projection (UMAP) colored by cell types (a) and species (b) and the scVI integration UMAP colored by cell types (c) and species (d). While PIANO has similar biological conservation scores to PCA and scVI and stronger batch correction than both, the human and mouse data are not well integrated in any integration, likely due to low counts per cell and strong species effects (e). Despite data quality issues, this integration is still useful for benchmarking runtime costs for training, where PIANO scales better than scVI to larger datasets, with up to 10 times faster training (f).

### PIANO outperforms scVI at mammalian whole brain integration

Another major challenge was to integrate a recently published whole human brain atlas^1^ with the whole mouse brain atlas^2^. Due to computational runtime limitations of scVI, we subset the data to up to 100 cells per highest resolution cluster in each atlas. We compared the biological conservation of brain regions and found cleanly separated clusters in the latent space embeddings (Figure 5a). Additionally, we found a good mixture of cells between human and mouse species across brain regions (Figure 5b). We also observed that the PIANO integration had comparable region separation to the scVI integration, and the PIANO batch correction performance steadily improved as the number of training epochs increased (Figure 5c). scVI regions were less separated (Figure 5d) and less mixed across species (Figure 5e). Although the PIANO integration benchmarking metric scores are lower than the previous examples, this is likely due to the complexity of the data and a limitation of using coarse brain regions as the "cell type" annotation. Moreover, the data, while not fully mixed across species like in the basal ganglia integration (Figure 3), is still fairly close together. The UMAPs for PCA can be found in Supplemental Figure 11, and benchmarking metrics scores can be found in Supplemental Table 16.

**Figure 5:**
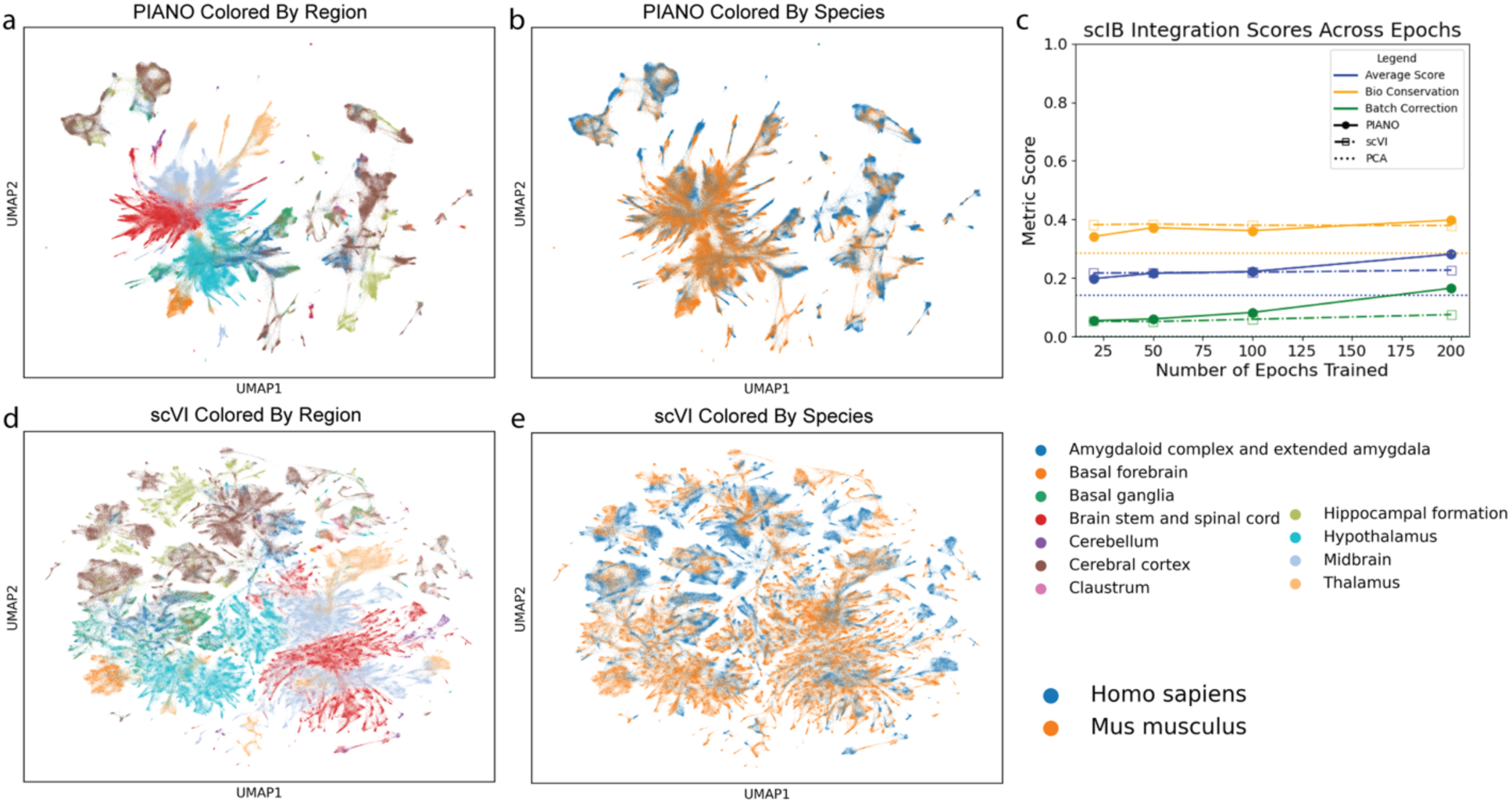
Whole mouse and human brain integration. PIANO enables robust integration of mouse and human whole brain atlases colored by region (a) and species (b). PIANO has stronger batch correction and stronger biological conservation of separation of brain regions than other methods (c). scVI integrations are shown colored by region (d), and species (e).

### PIANO enables multi-modal integration with spatial transcriptomics

Integration across data modalities empowers biological discovery. Spatial transcriptomics technologies, such as multiplexed error-robust fluorescence in situ hybridization (MERFISH)^29^, enable identifying exact physical locations of cells in a slice of tissue, which is not known in single cell RNA transcriptomics data; however, this spatial resolution comes at the cost of capturing fewer genes than single cell data. The Broad Institute created a whole mouse brain spatial atlas containing a complete spatial transcriptomics series of four mouse brains^30^; due to having more genes, we decided to use the Broad Institute’s MERFISH data with 1,122 genes shared across panels^30^ instead of the Allen Institute’s MERFISH data with a 500 gene panel^2^ for demonstrating spatial integration.

To obtain cell type labels, the Broad Institute spatial atlas was originally integrated with the Allen Institute’s whole mouse brain single cell atlas^2^ using Seurat (CCA)^30,31^. Given PIANO and scVI’s superior performance to Seurat for data integration, we revisited integrating the Broad Institute’s whole mouse brain spatial transcriptomics atlas with the Allen Institute’s whole mouse brain single cell transcriptomics atlas. We first subset the genes in the single cell transcriptomics data and spatial transcriptomics data to the 1,122 genes shared in all datasets^2,30^; we then subset the cells in each datasets to up to 6400 cells per Subclass to obtain a balanced dataset for integration. We found good mixing between single cell and spatial modalities (Figure 6a) and good separation of cell type Classes (Figure 6b). Since the Allen Institute whole mouse brain taxonomy used hierarchical clustering for cell types, we demonstrated an illustrative example of hierarchical reintegration for the cells from Class 13 CNU-HYa Glut at the Subclass level (Figure 6c). Since the training time for PIANO is approximately linear with respect to the number of cells, PIANO can perform hierarchical integration with a linear runtime cost with respect to the number of levels in the hierarchy. This enables efficient and comprehensive analyses of large atlases at multiple cell type resolutions. In contrast, scVI had less mixing across modalities (Figure 6d), less separation of classes (Figure 6e), and less separation of subclasses (Figure 6f). Moreover, PIANO had better mixing of modalities for subclasses (Figure 6g) than scVI (Figure 6h). Again, we observed superior biological conservation and batch correction compared to scVI for both the whole brain integrations (Figure 6i) and Class 13 CNU-HYa Glut integration (Figure 6j). UMAPs for PCA can be found in Supplemental Figure 12, and benchmarking scores can be found in supplemental tables 17-18.

**Figure 6:**
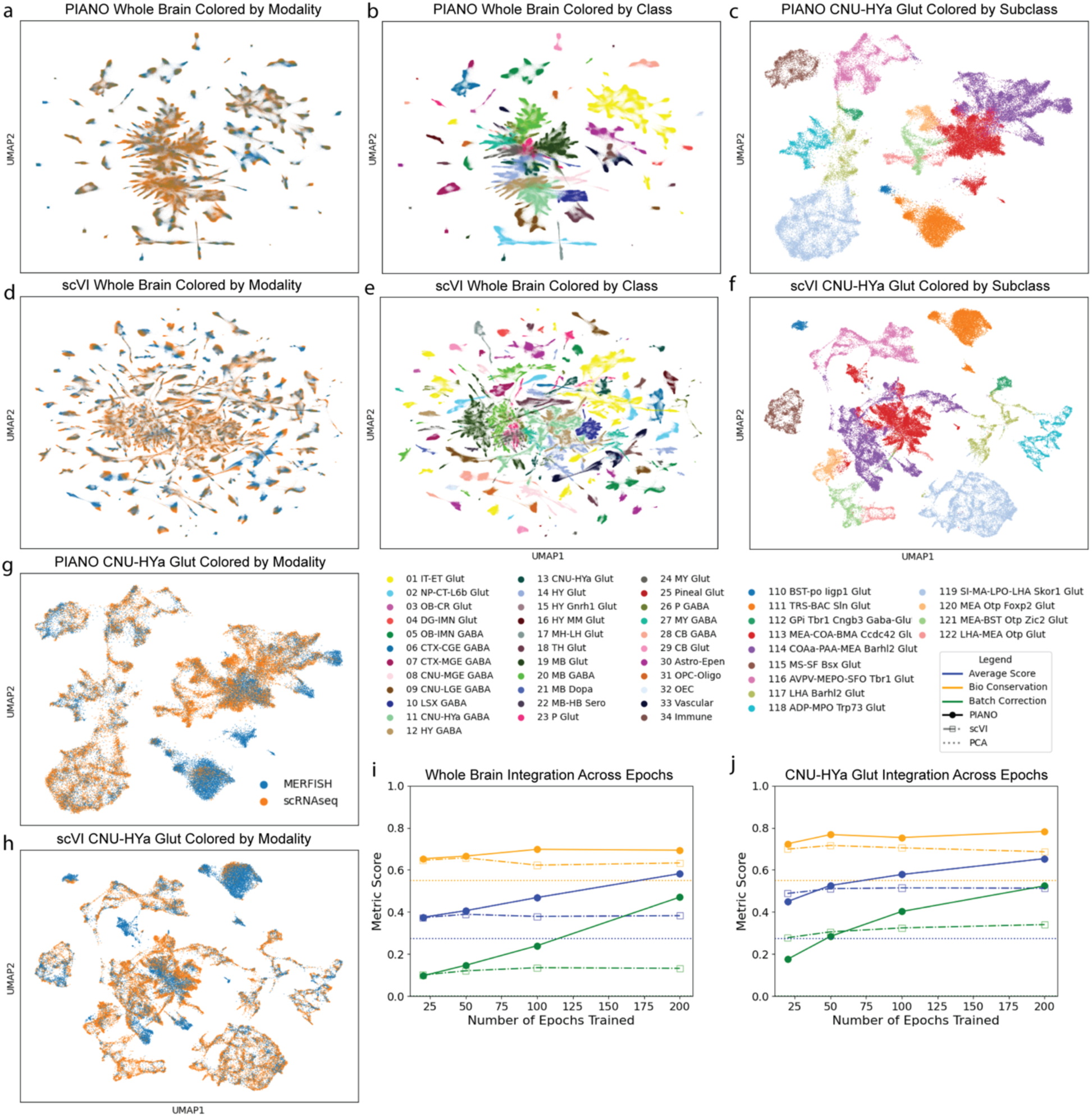
Whole mouse brain spatial integration. PIANO integrates mouse whole brain 10X single cell and MERFISH spatial transcriptomic atlases colored by sequencing modality (a) and Class (b), and Class 13 CNU-HYa Glut is shown colored by Subclass (c). scVI integrations are shown colored by sequencing modality (d) and Class (e), and Class 13 CNU-HYa Glut is shown colored by Subclass (f). Class 13 CNU-HYa Glut integrations are shown colored by modality for PIANO (g) and scVI (h). PIANO outperforms scVI for whole brain integration performance (i) and for Class 13 CNU-HYa Glut integration performance (j).

### PIANO enables integration across species and developmental timepoints

Developmental datasets pose difficult challenges for integration tasks, as cell type identities are especially dynamic during maturation^26^. This challenge is further exacerbated when attempting to integrate across species, since developmental timepoints may not be aligned. In a recent study, we compared astrocyte development across embryonic and postnatal timepoints in mice and marmosets, but were not able to successfully integrate the species datasets together using existing methods^26^.

Using PIANO, we obtained robust integrations of the complete dataset, including neurons and non-neuronal cells^26^ (Figure 7). We found strong integration with good mixing across species (Figure 7a), separation of cell types (Figure 7b) and superclusters (Figure 7c). In contrast, scVI had less mixing across species (Figure 7d), and similar separation of cell types (Figure 7e) and superclusters (Figure 7f). Additionally, we found separation along certain developmental time points, such as cells in the embryonic stage distinctly separated from the clusters in the later time points (Figure 7g), which were mixed together in the previous single species integrations^26^. Moreover, we preserved distinct region differences observed in both neurons and glia (Figure 7h), recapitulating region specificity of glial cell types as described in the original atlas manuscript^26^. Although scVI similarly captured these differences (Figure 7i-j), it struggled with mixing data across species (Figure 7k). Furthermore, the original manuscript also identified two species-specific neuronal superclusters: an embryonic mouse type ’Inh Str Immature’, and a marmoset thalamic GABAergic type ’Inh Thal MB-der’; both species-specific cell types are preserved in PIANO’s integration (Figure 7a-c,g-h). These findings show that PIANO can perform highly-robust cross-species integration comparisons between evolutionary distant species, while still preserving biological signals attributable to regional and developmental sources. UMAPs for PCA can be found in Supplemental Figure 13, and benchmarking metrics are in Supplemental Table 19.

**Figure 7:**
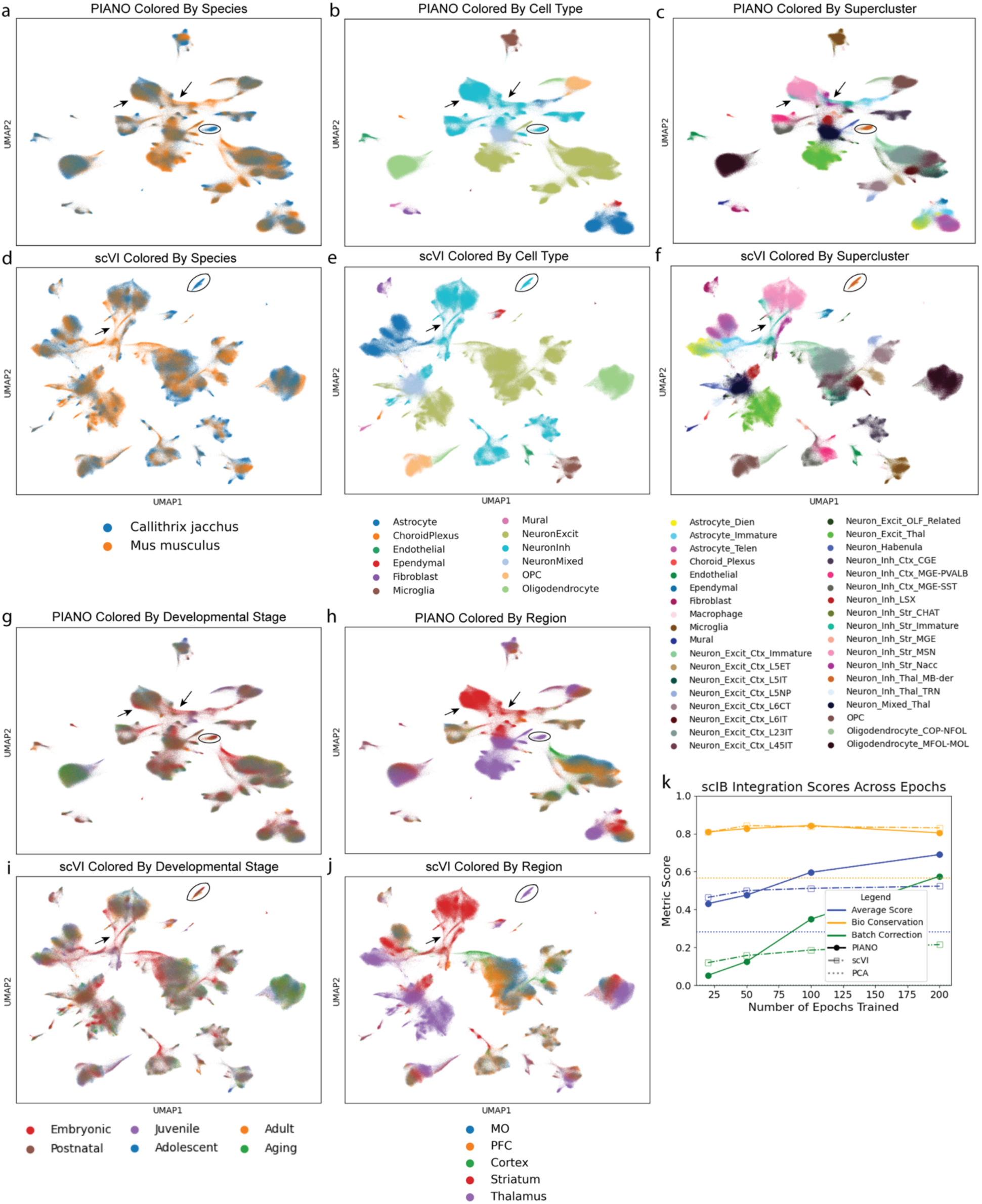
Developmental mouse and marmoset brain integration. PIANO enables robust integration of mouse and marmoset cell types across tissues and developmental stages colored by species (a), cell type (b), supercluster (c), with scVI integrations colored by species (d), cell type (e), supercluster (f). PIANO integrations are colored by developmental stage (g), and brain region (h), and scVI integrations are colored by developmental stage (i), and brain region (j). PIANO has superior batch correction and similar biological conservation to scVI (k). A marmoset-specific thalamic supercluster is circled, and a mouse-specific embryonic supercluster is indicated by the arrows (a-j).

## DISCUSSION

### PIANO enables fast and robust integrations

We introduce PIANO, a powerful, scalable, and efficient framework for integrating single cell multi-omics data. We showed that PIANO enables 1) fast integrations that require a small fraction of the runtime cost of existing tools and 2) produces high-quality latent space representations that both preserve desired biological features and integrate data between multiple batches. We demonstrate applications of our integration framework across a wide variety of datasets over multiple species, tissues, sequencing platforms, and developmental timepoints. The broader impacts of our method include greatly expanding the accessibility of powerful GPU-based integration tools by reducing the runtime cost by an order of magnitude while producing superior or comparable integration results. We also dramatically speed up training with our light-weight codebase, enabling atlas-scale integrations in less than an hour.

### Model parameter selection

The default PIANO model architecture with 3 hidden layers, 256 hidden nodes, and 32 latent dimensions worked well for all integrations with 200 training epochs. The most important PIANO hyperparameter to adjust is the max beta hyperparameter, where a higher max value for beta results in stronger batch correction in exchange for some biological conservation of cell types. For most of our integrations, we used our default max value of 1.0 after 400 training epochs (see Methods). In situations where the default PIANO beta value results in over-mixing across batches, we recommend reducing the max value. This is exemplified by the withheld D1 MSN primate basal ganglia integration, where reducing the max value from 1.0 to 0.5 recaptured the human-specific perturbed cell type (Supplemental Figure 9). PIANO trains quickly using code compilation and GPU acceleration, which makes hyperparameter sweeps computationally tractable. We also recommend first identifying cell types within each species before performing cross species integration to confirm that biological signal present in the single species analysis is not lost or distorted upon integration.

### Gene selection

Previous studies have found that some functional gene sets, such as transcription factor genes, are able to more accurately predict cell types than other gene sets, such as cell adhesion molecule encoding genes^2^. We found for integration across species, all functional gene sets performed similarly, though larger numbers of genes generally had better performance (Figure 3j). This is consistent with our comparison over the number of highly variable genes, where more genes used resulted in slightly higher biological conservation in exchange for small decreases in batch correction (Figure 3h), and our comparison with increasing numbers of random genes shows a similar pattern (Figure 3i). Although functional gene sets may be informative, the number of genes used for integration is also an important consideration.

### Data quality influences integration results

One notable observation is the variation in the quality of integrations between various atlases. In particular, the mouse and marmoset developmental integration from Schroeder et al.^26^ produced very clean integrations with both exceptionally high biological conservation and batch correction scores (Figure 6), whereas the mouse and human cross-tissue integration between the MCA and HCL datasets had far lower integration performances (Figure 4). While this may be impacted by sampling different tissue types, there is also a stark difference in the sequencing depth and ultimate UMI yield between the datasets. Schroeder et al. targeted 40,000 sense reads per nucleus^26^, whereas the HCL obtained a substantially fewer 3000 reads per cell^28^. Moreover, Schroeder et al. used the 10x 3’ v3 chemistry^26^, whereas the HCL used a Microwell-seq platform^28^. As a result, the mean number of unique molecular identifiers (UMIs) per nuclei in each batch had a mean of 6,085 UMIs per nuclei across 144 batches in Schroeder et al.^26^, whereas the HCL data had a mean of less than 934 UMIs per cell in each tissue across 112 tissue samples^28^. For the whole brain integrations, the Allen Institute’s whole mouse brain atlas achieved an average of 54,379 reads per cell for the 10Xv2 data and 83,190 reads per cell for the 10Xv3 data^2^, and the whole human brain atlas achieved approximately 100,000 reads per nuclei^1^. Note these datasets count both sense and anti-sense reads, while the Schroeder et al. study only counted sense reads. As anti-sense reads can make up to 40% of reads, these datasets are all generally comparable in sequencing depth.

Dramatic differences in dataset depth can contribute to the large discrepancies in the quality of integration results. This reinforces the need for high-quality sequencing data to distinguish between evolutionary divergences across species with confounding noise artifacts. Just as Seurat was inspired by pointillism painting, the name of the our method, PIANO, serves as a metaphor between integrating complex atlases to performing piano concertos, such as Rachmaninoff’s Piano Concerto No. 2 in C minor^32,33^. Just as the quality of the performance depends on individual contributions of the pianist, conductor, and orchestra, the success of cross-species integrations relies on having large quantities of high-quality data. While the PIANO integrations increased batch correction across epochs at a higher rate than scVI across most datasets, this performance improvement was more apparent in datasets in which a large number of cells were available for each cell type when training the model (Methods). This further highlights the importance of high-quality training data for robust integration performance.

### Limitations

One limitation of our model is that it relies on modern GPU acceleration to make the most of its performance improvements. While we tested our model for both x86 and ARM CPU architectures, the best performance improvements come with its compiled mode, which requires the use of a GPU that can support torch.compile optimizations. Moreover, these compilations introduce up to two minutes of additional compilation time overhead costs, which make up a larger proportion of runtime for smaller datasets. These hardware requirements can be mitigated using institutional high-performance computing clusters (HPCs) or cloud computing platforms.

Finally, as with most machine learning methods, the quality of the results depends on the quality of the input data. Using more high-quality data not only results in more substantial integration and runtime performance improvements compared to other integration methods, but also further enriches the training of the model. Moreover, with single cell integration, there still exists a fundamental tradeoff between batch correction and biological conservation^34^.

### Related work

The NB-GLM modeling used in the VAE decoder was inspired by the NB-GLM used by scTransform for normalizing scRNAseq data^35^. Other commonly used integration tools include Seurat (CCA)^6^, Harmony^5^, scVI^9^, and others. Additionally, some integration methods attempt to use a semi-supervised approach, such as scANVI^36^. However, these methods rely on the robustness of the assigned cell type labels and may bias results against novel biology or rarer cell types, and do not model relationships between similar cell types. Moreover, using a semi-supervised approach may introduce training instability and longer runtimes.

## METHODS

### PIANO model architecture

PIANO uses approximate Bayesian inference to train a modified variational autoencoder (VAE)^8^ using an encoder-decoder architecture. The encoder, or recognition model, assumes an isotropic Gaussian prior and learns a probabilistic, posterior latent space representation for the observed input data X (Figure 1). This latent space uses an independent mean and standard deviation for each dimension (Figure 1), which enables sampling via a reparameterization trick^8^. The decoder, or generative model, samples a latent variable, Z, which is combined with the covariates, C, as its inputs to learn the parameters for a negative binomial generalized linear model (NB-GLM) that maximizes the likelihood of the data.

For a given cell, each gene is modeled using a negative binomial (NB) distribution: X_gene_ ∼ NB(ⲝ_gene_, 𝜓_gene_). The NB distribution reflects the discrete nature of mRNA transcripts^37^. Specifically, the NB distribution models the number of successes obtained before r failures for a sequence of independent and identically distributed (IID) Bernoulli trials, where each trial is analogous to whether a cell expresses a specific mRNA transcript. The NB distribution for each gene is parameterized using a generalized linear model (GLM) for ⲝ_gene_, the overdispersion (i.e., the number of failures before stopping), and 𝜓_gene_, the log-odds of success.

Specifically, we model the overdispersion per gene with 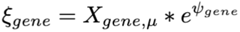, where 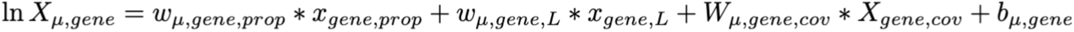 is the predicted log expression of a gene from the decoder. Here, X_gene,prop_ is the output of the decoder after a softmax transformation, which produces the proportion of transcripts for each gene for a given cell. This proportion is scaled by the library size (total number of transcripts) for each cell, X_gene,L_, which influences the mean gene expression^35^. X_gene,cov_ is a vector of the covariates for a given cell, which include categorical covariates (one-hot encoded) and continuous covariates (z-scored). W_µ,gene,cov_ is a vector of weights for the covariates, w_µ,gene,prop_ is a scalar weight for the gene proportion, w_µ,gene,L_ is a scalar weight for the library size, and the bias term is given by b_µ,gene_. This differs from scVI’s implementation, which does not include a bias term and uses weights for only the primary batch key^9^ as opposed to multiple covariates: 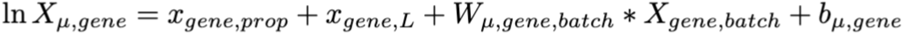

PIANO also models the other NB parameter, the log-odds for capturing a transcript, using 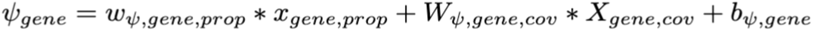. Here, w_𝜓,gene,prop_ is the scalar weight for the gene proportion, W_𝜓,gene,cov_ is the vector of weights for the covariates, and b_𝜓,gene_ is the bias term. We assume that the library size should only impact the mean expression of genes, and therefore, do not include it to model the log-odds. scVI uses a mean-overdispersion parameterization^9^, which does not model log-odds.

The likelihood of the data under the PIANO NB-GLM is given by:

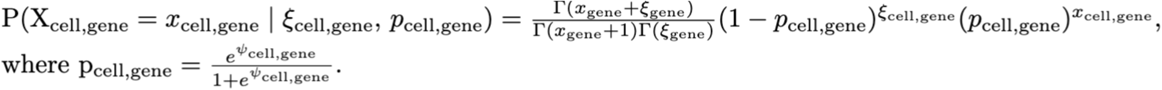

### PIANO evidence lower bound objective

PIANO is trained by minimizing an evidence lower bound objective (ELBO), which is a lower bound on the log-probability of the observed gene expression data given the covariates. The ELBO is the sum of a data fidelity term, given by the average log-likelihood of the data under the NB observation model, and the negative Kullback-Leibler divergence (KLD) between the approximate posterior over Z from the recognition model and the isotropic Gaussian prior. The total ELBO is the sum of ELBOs across cells. The ELBO can be derived as follows:

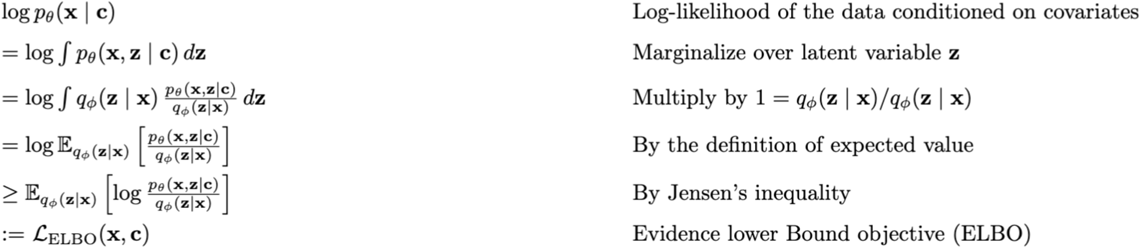

The ELBO can be written as an expected log-likelihood and a regularizing KLD term^8^.

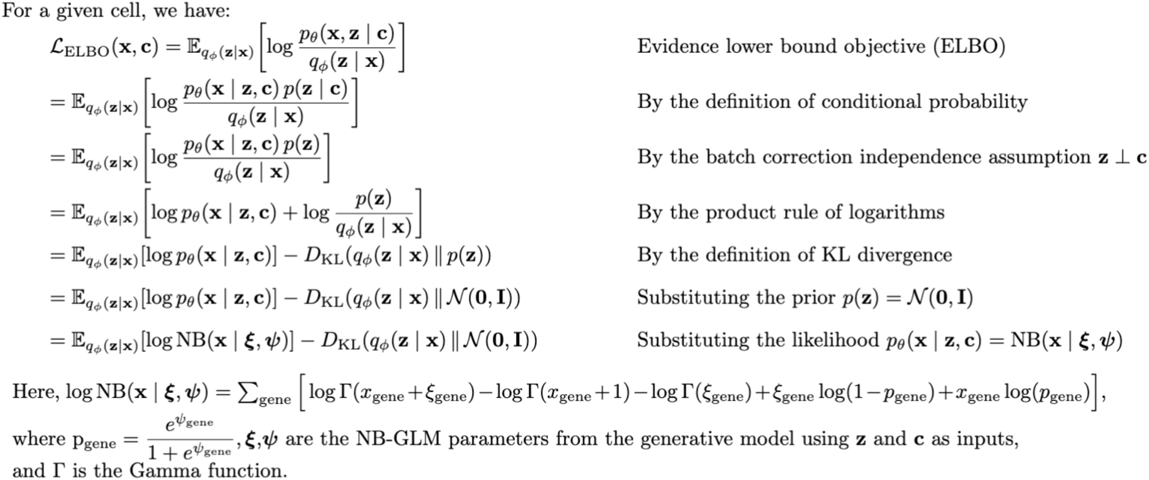

The KLD term is calculated analytically between the encoder’s latent space distribution, which is a diagonal Gaussian, and an isotropic Gaussian prior distribution^8^. The log-likelihood is calculated using the log-likelihood of the original data under the NB-GLM with the corresponding covariates. By sampling from the latent space representation, the generative model produces an expectation over the log-likelihood of the data. This VAE model architecture enables PIANO to train in an unsupervised manner.

### Implementation details

The encoder, or recognition model, learns an approximate posterior distribution over the latent P(Z|X) (e.g., 32 latent dimensions) given a vector of gene counts (e.g., 4096 highly variable genes); specifically, the encoder outputs a probabilistic posterior latent representation for each cell using an independent mean and standard deviation for each latent dimension. Using the reparameterization trick^8^, a z-score is sampled outside of the computational graph to produce the posterior latent representation P(Z|X) from the mean, standard deviation, and z-score. This latent representation is concatenated with covariates as the input to the decoder.

Given a latent vector and a set of covariates (e.g., one-hot encoding for 3 primate species), the decoder parametrizes P(X|Z,C) (i.e., a vector for 4096 genes), which is the proportion of gene counts; specifically, the decoder uses softmax to obtain gene proportions that sum to one. These proportions are then scaled by the library size (total counts) of a given cell to obtain the means of the negative binomial distributions over the genes. Compressing the data through the VAE bottleneck helps the latent space representation capture only the most important biological information for reconstructing the gene proportions. Feeding the covariates into only the decoder helps the model remove batch effects from the latent space while still modeling the original gene counts, which requires information from the covariates.

Both the encoder and decoder are implemented in PyTorch using a multi-layer perceptron (MLP), or feedforward neural network, following the pattern of feed-forward linear layer, batch normalization, rectified linear unit (ReLU), and dropout. PIANO uses a default network architecture with 3 layers, 256 hidden nodes per linear layer, and 32 latent dimensions. PIANO uses default parameters with 1e-5 for the batch normalization epsilon, 1e-1 for batchnorm momentum, and 0.1 for dropout rate. PIANO uses an AdamW optimizer with a default learning rate of 2e-4, mini-batch size of 128 cells, and no weight decay. For training stability, datasets with normalized total transcript counts in the hundreds of thousands per cell are first mean-scaled to 10,000 transcripts per cell within each batch; the gene expression data are also log1p-transformed before being used as input for the encoder. For each training step, we use a mini-batch update with 128 randomly sampled cells without replacement.

The two most important parameters for PIANO are the number of training epochs and the hyperparameter beta, which govern the tradeoff between biological conservation and batch correction. The number of training epochs increases the training time linearly. Although training costs are not an issue with PIANO, due to its torch.compile GPU acceleration, training the model for too many epochs may result in over-correction for batch effects and risk losing separation of cell types. To mitigate this potential issue, PIANO is designed with easily accessible model checkpoints at fixed training intervals; a visual inspection of the latent space UMAP at each training interval reveals when the model has achieved an acceptable balance between batch correction and biological conservation. Using GPU acceleration from the RAPIDS AI library, UMAP can be computed within seconds even for large datasets. This enables visual inspections to determine whether PIANO has finished training, which helps avoid over-correction of batch effects.

In practice, VAEs are trained using beta-annealing. We scale linearly with every step, where the beta weight is 1.0 after 400 training epochs have completed. Specifically, we have 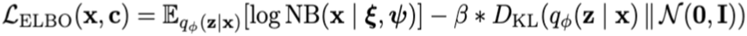.

To mitigate KLD vanishing, PIANO uses a KLD beta-annealing schedule^38^ that linearly increases the KLD weight after each mini-batch update. The KLD term is weighted using a beta hyperparameter that increases after each training step; a larger weight generally results in stronger batch correction. PIANO uses a linear KLD annealing schedule starting from an initial weight of 0.0 to its maximum weight after 400 epochs, incrementing after every training step^38^. Although 400 training epochs are usually not necessary, this large upper bound makes the default schedule sufficient for most applications. The max KLD weight is the most important hyperparameter to tune for balancing batch correction with biological conservation of cell types.

We found a max beta of 1.0 to be effective for most datasets. For the lung integration, we used a weaker max beta of 0.5. For most integrations, we used our default model parameters, architecture, and 200 max training epochs (Methods). Additionally, we found our default learning rate of 2e-4 to be effective for the larger datasets, but reduced the learning rate to 1e-4 and used a weight decay of 1e-6 to avoid overfitting the smaller datasets (pancreas, lung, immune, and tabula muris) that had fewer than 50,000 cells; these datasets may have had insufficient training samples for the models to train properly. Moreover, we found our default model architecture design with 3 hidden layers, 256 hidden nodes per hidden layer, and 32 latent dimensions effective on all datasets (Methods). In general, it is sufficient to run PIANO with the default parameters, using model checkpoints to determine if using fewer training epochs are sufficient. If over-correcting for batch effects, PIANO can be run with a lower beta (i.e. 0.5 instead of 1).

### Code compilation

PIANO is implemented in PyTorch and achieves faster training through backpropagation compared to existing VAE methods using the torch.compile function. Using a custom training loop, the entire training step is compiled, including both forward and backward passes. This loop also enables model checkpointing without the use of PyTorch Lightning callbacks. A PyTorch DataLoader with a custom Sampler (e.g. GPUBatchSampler) is used to efficiently extract mini-batches from the data, which is stored in a custom PyTorch Dataset class (AnnDataset). This AnnDataset class stores the raw counts and covariates in an augmented data matrix, which supports dense GPU memory mode, a custom sparse GPU memory mode, and CPU memory modes; this allows flexibility depending on the size of the data and availability of GPU memory. These code optimizations enable PIANO to train substantially faster than other VAE methods.

### Preprocessing

All integration examples used publicly available and preprocessed datasets; as such, no new quality control thresholds were used. For cross-species comparisons, we subset the data to the human one-to-one orthologs from Ensemble version 114^22^. For the primate basal ganglia integration gene sets, one-to-one ortholog transcription factors were selected from AnimalTFDB4^24^ and one-to-one ortholog functional gene sets were queried from UniProtKB^23,39^. Scanpy’s sc.pp.highly_variable_genes was used to select 4096 genes for most integrations, using the ’seurat_v3’ flavor and the most prominent source of variation for the batch key^31,40^. For the single species dataset integrations in Figure 1, the batch keys used were sequencing platform for pancreas, donor for lung, dataset batch for immune, donor for tabula muris, and donor for heart atlas. For each of the cross-species integrations, species was used as the batch key. For the whole mouse brain spatial integration, all 1,122 genes were used.

For most datasets, the raw gene counts were used as input. For datasets with substantially larger magnitudes for the raw data (i.e. due to normalizing differences in the sequencing data acquisition), each dataset was scaled to a mean of 10k. These datasets include ’Villani’ for the immune integration, the ’fluidigmc1’, ’smarter’, and ’smartseq2’ datasets for the pancreas integration, and all donors for the tabula muris integration.

To balance the number of cells between datasets, we used 6400 cells per group for each species in the primate basal ganglia, up to 2^18^ (262,144) cells per species (mouse, human) for each of the Mouse Cell Atlas (MCA) and Human Cell Landscape (HCL), 100 cells per highest resolution cluster for each of the mouse and human whole brain datasets, 6,400 cells per Subclass for each modality (scRNAseq, spatial) in the whole mouse brain integration, and 2^16^ (65,536) cells per developmental timepoint for each species (mouse and marmoset) for the developmental integration.

For principal component analysis (PCA), 30 principal components were used in each integration example. For UMAP visualization, the Scanpy default 15 neighbors were used to compute the neighborhood graph. For Seurat, the k.weight was set to 50 (as opposed to the default 100 to enable integrations on batches with fewer cells). For Harmony, default parameters were used with max 20 iterations. For scVI, the dispersion for its zero-inflated negative binomial distribution (as described in their original manuscript) was set to ’gene-batch’, and all other parameters were set to defaults besides the model architecture, which was set to the same 3 hidden layers, 256 hidden nodes per hidden layer, and 32 latent dimensions used by PIANO for benchmarking comparisons.

### Benchmarking

Integration embeddings were evaluated by biological conservation metrics, which measure separation of cell type clusters, and batch correction metrics, which measure how mixed cells are between different batches. The biological conservation metrics used were normalized mutual information (NMI) and adjusted Rand index (ARI). The batch correction metrics used were the k-nearest-neighbors batch effect test (KBET) and inverse local Simpson’s index (iLISI)^7^. It is well understood that a good integration should maintain separation of cell type clusters while mixing data between batches without over correction^7^.

The scib_metrics package was used to run integration benchmarking with commonly used metrics^7^ The average of the Leiden Adjusted Rand Index (ARI) and Leiden Normalized Mutual Information (NMI) scores was used to measure biological conservation; both metrics use de novo Leiden clustering on the latent space representation of the data and compare the clusters to the ground truth labels^7^. The average of the K-nearest neighbors Batch Effect Test (KBET) and integration Local Inverse Simpson’s Index (iLISI) scores were used to quantify batch correction; these metrics compare the number of cells from different batches in the neighbors of a given cell^7^.

## Supporting information

Key Resources Table

## Code availability

The PIANO source code is publicly available at https://github.com/NingWang1729/piano with an installable package at https://pypi.org/project/piano-integration. A publicly available Code Ocean capsule demonstrating the human and mouse whole brain integration is available at https://codeocean.allenneuraldynamics.org/capsule/4212260/tree/v2.

## Data availability

All datasets used were already preprocessed and publicly available. Download links can be found in the key resources table.

## Author Contributions

N.W., C.L., J.W.P., and F.M.K. designed the method. N.W. and D.T. implemented the method. N.W., H.F., and V.N. benchmarked other methods. N.W., H.F., V.N., N.S., C.L., D.Y., Z.Y., C.C., M.D., S.D., L.C., and J.S. performed data collection, preprocessing, and analysis. N.W., C.L., J.W.P., and F.M.K. wrote the manuscript with input from all authors.

## Acknowledgements

This publication was supported by and coordinated through the Brain Initiative Cell Atlas Network (BICAN). This work was supported by National Institutes of Health UM1MH130981, DP2MH140136, and the Klingenstein–Simons Fellowship (F.M.K.). N.W. was supported by The McDonnell Fellows in Neuroscience at Princeton University and National Institutes of Health 5T32MH065214. This work was completed in part at the Princeton Open Hackathon, part of the Open Hackathons program. The authors would like to acknowledge OpenACC-Standard.org for their support. The authors would like to acknowledge the Allen Institute for Neural Dynamics for their computational support with Code Ocean. We wish to thank the Allen Institute founder, Paul G. Allen, for his vision, encouragement, and support.

