## Supplemental Figures for "Fast Optimization of Robust Transcriptomics Embeddings using Probabilistic Inference Autoencoder Networks for multi-Omics"

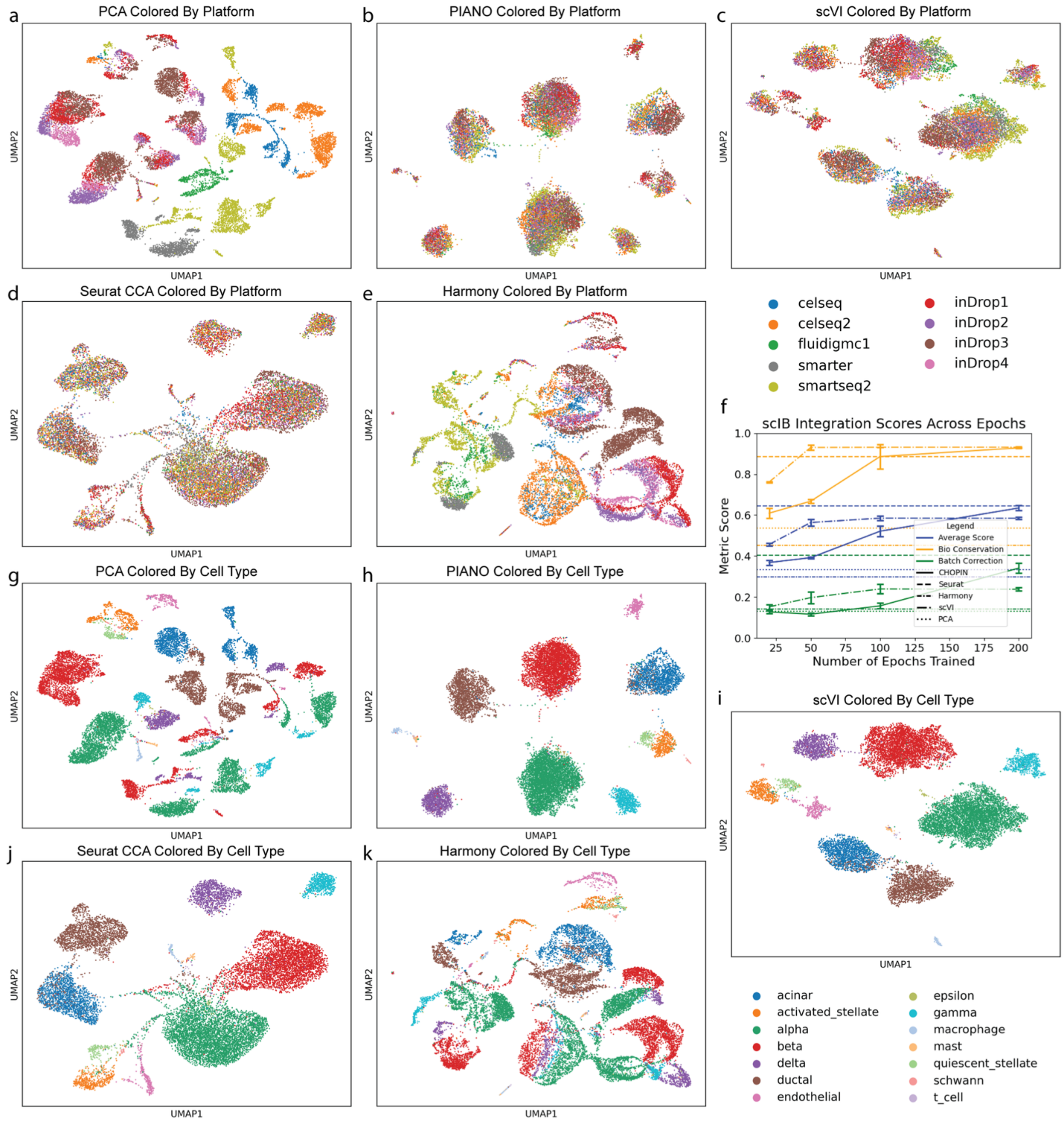


Figure S1: Pancreas integration using multiple methods. Uniform manifold approximation and projections (UMAPs) of pancreas data colored by platform for PCA (a), PIANO (b), scVI (c), Seurat (d), and Harmony (e). Integration performances are shown (f). UMAPs of pancreas data colored by cell type for PCA (g), PIANO (h), scVI (i), Seurat (j), and Harmony (k).

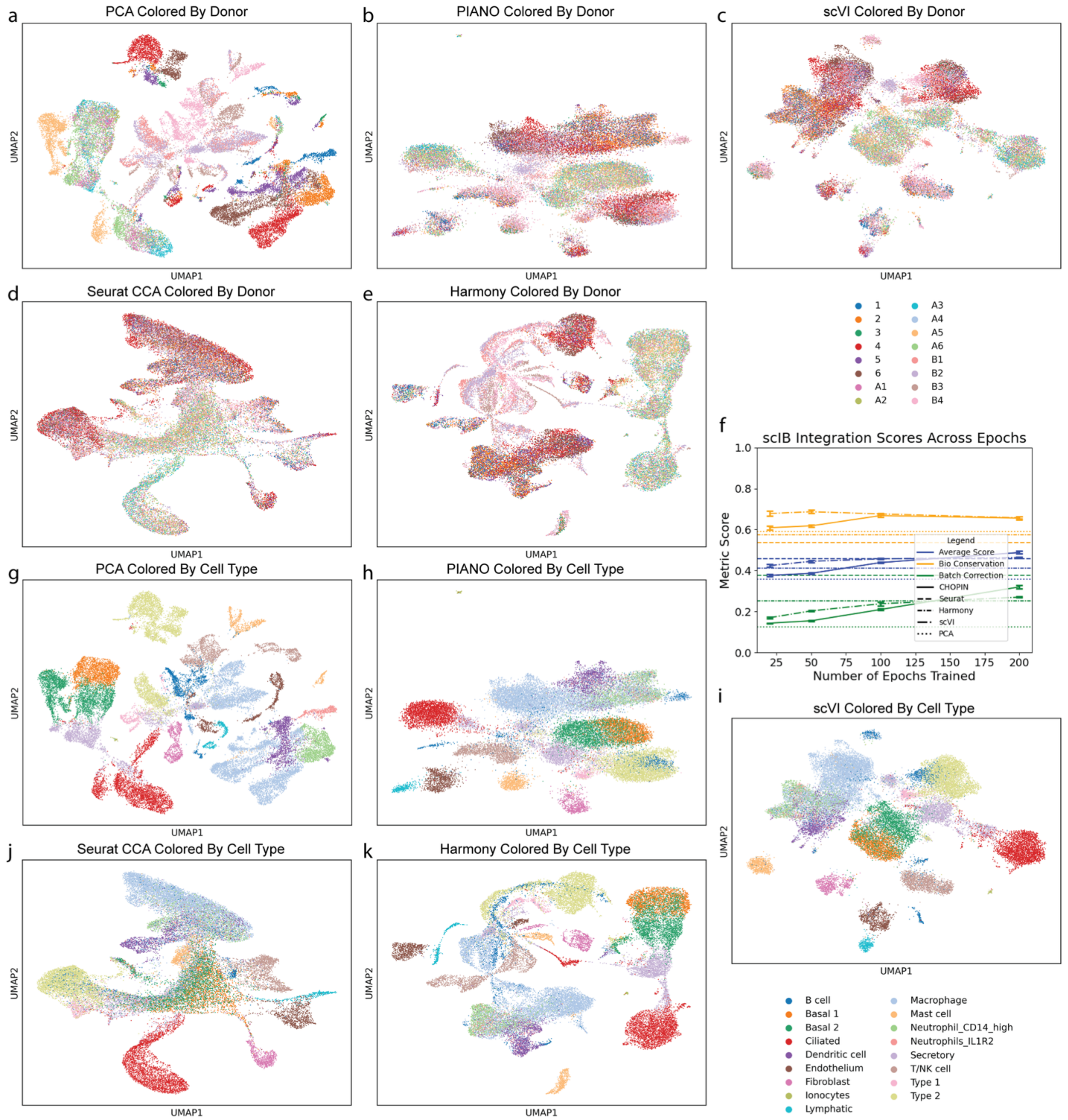


Figure S2: Lung integration using multiple methods. Uniform manifold approximation and projections (UMAPs) of lung data colored by platform for PCA (a), PIANO (b), scVI (c), Seurat (d), and Harmony (e). Integration performances are shown (f). UMAPs of lung data colored by cell type for PCA (g), PIANO (h), scVI (i), Seurat (j), and Harmony (k).

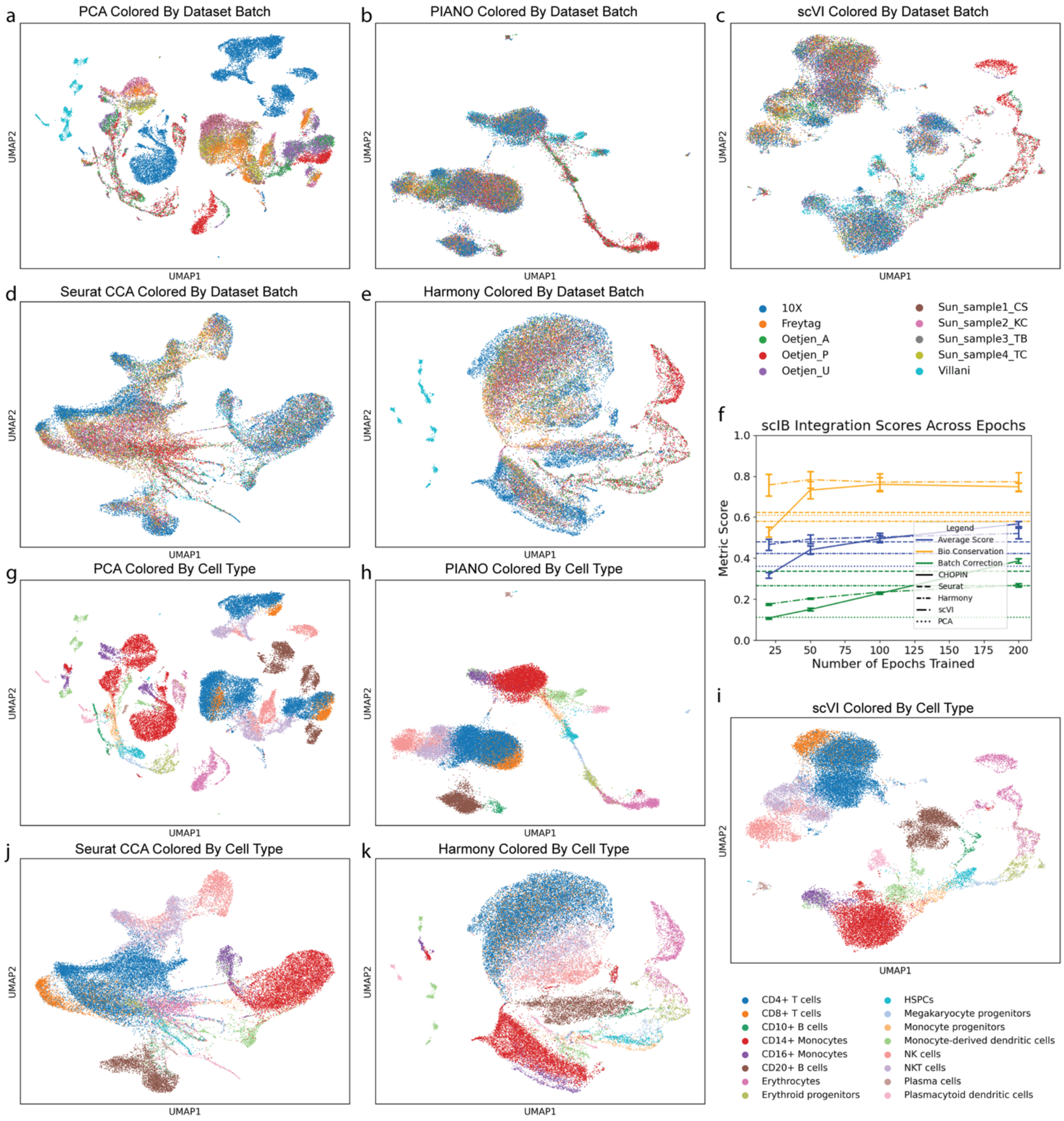


Figure S3: Immune integration using multiple methods. Uniform manifold approximation and projections (UMAPs) of immune data colored by platform for PCA (a), PIANO (b), scVI (c), Seurat (d), and Harmony (e). Integration performances are shown (f). UMAPs of immune data colored by cell type for PCA (g), PIANO (h), scVI (i), Seurat (j), and Harmony (k).

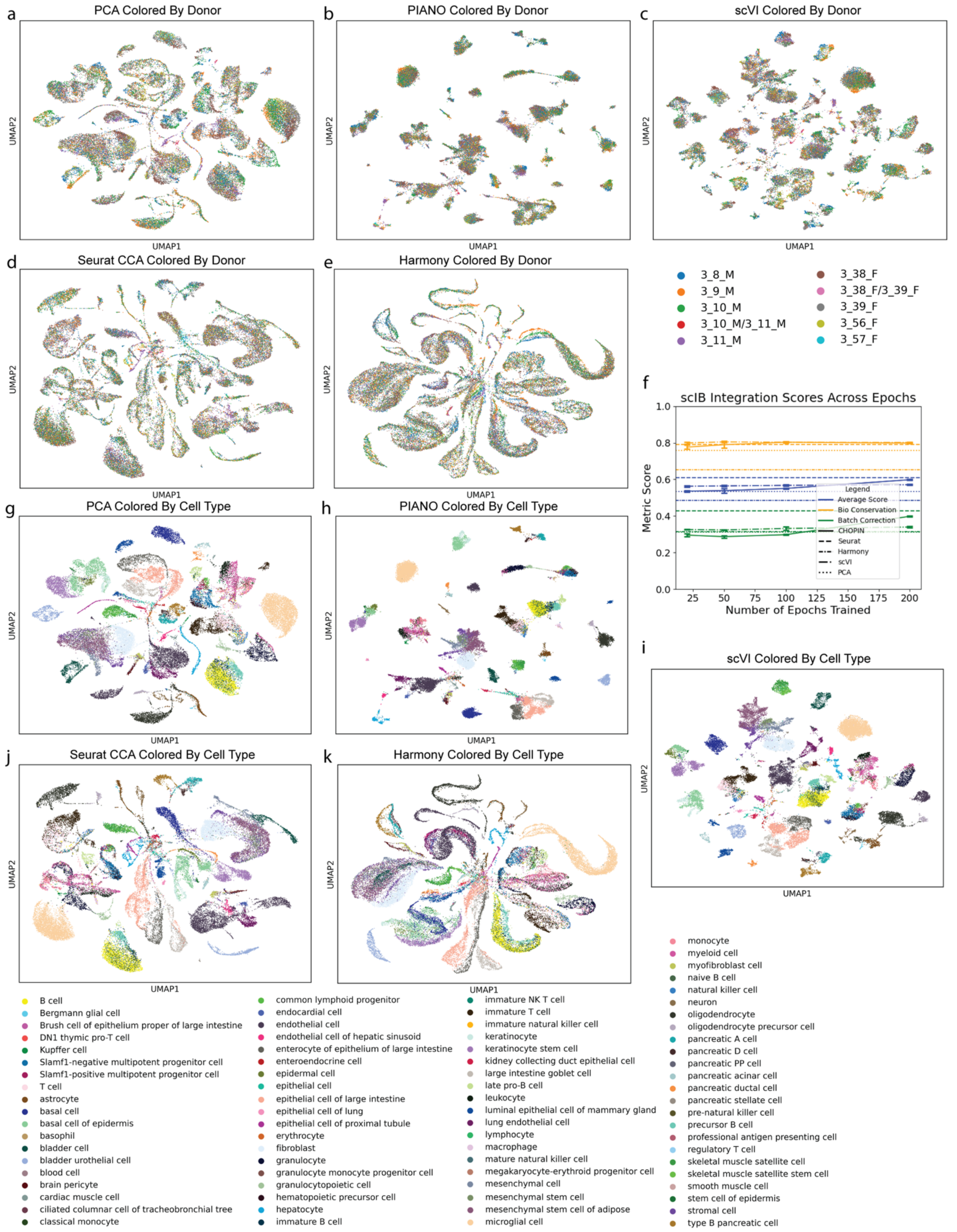


Figure S4: Tabula Muris integration using multiple methods. Uniform manifold approximation and projections (UMAPs) of tabula muris data colored by platform for PCA (a), PIANO (b), scVI (c), Seurat (d), and Harmony (e). Integration performances are shown (f). UMAPs of tabula muris data colored by cell type for PCA (g), PIANO (h), scVI (i), Seurat (j), and Harmony (k).

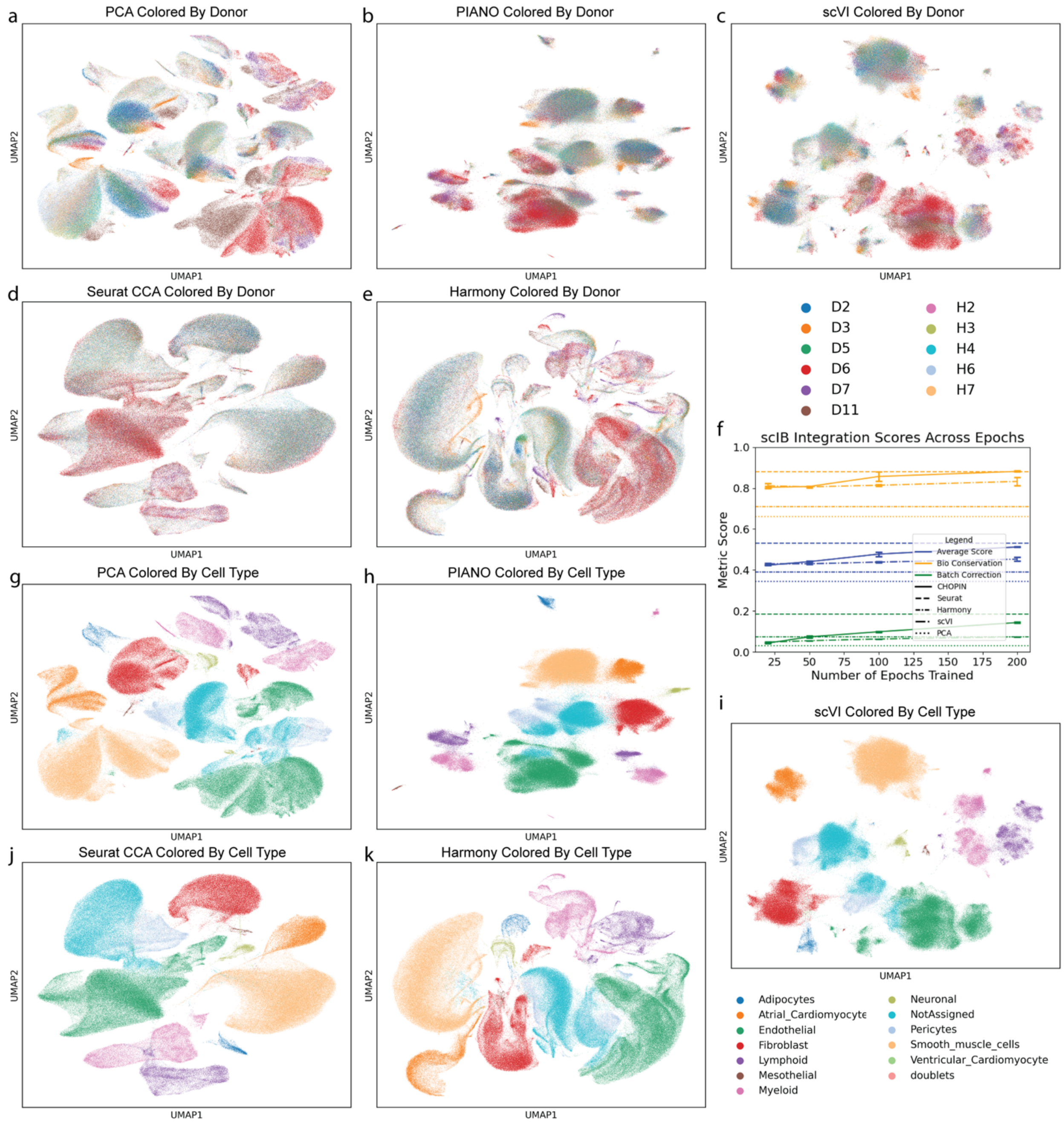


Figure S5: Heart integration using multiple methods. Uniform manifold approximation and projections (UMAPs) of heart data colored by platform for PCA (a), PIANO (b), scVI (c), Seurat (d), and Harmony (e). Integration performances are shown (f). UMAPs of heart data colored by cell type for PCA (g), PIANO (h), scVI (i), Seurat (j), and Harmony (k).

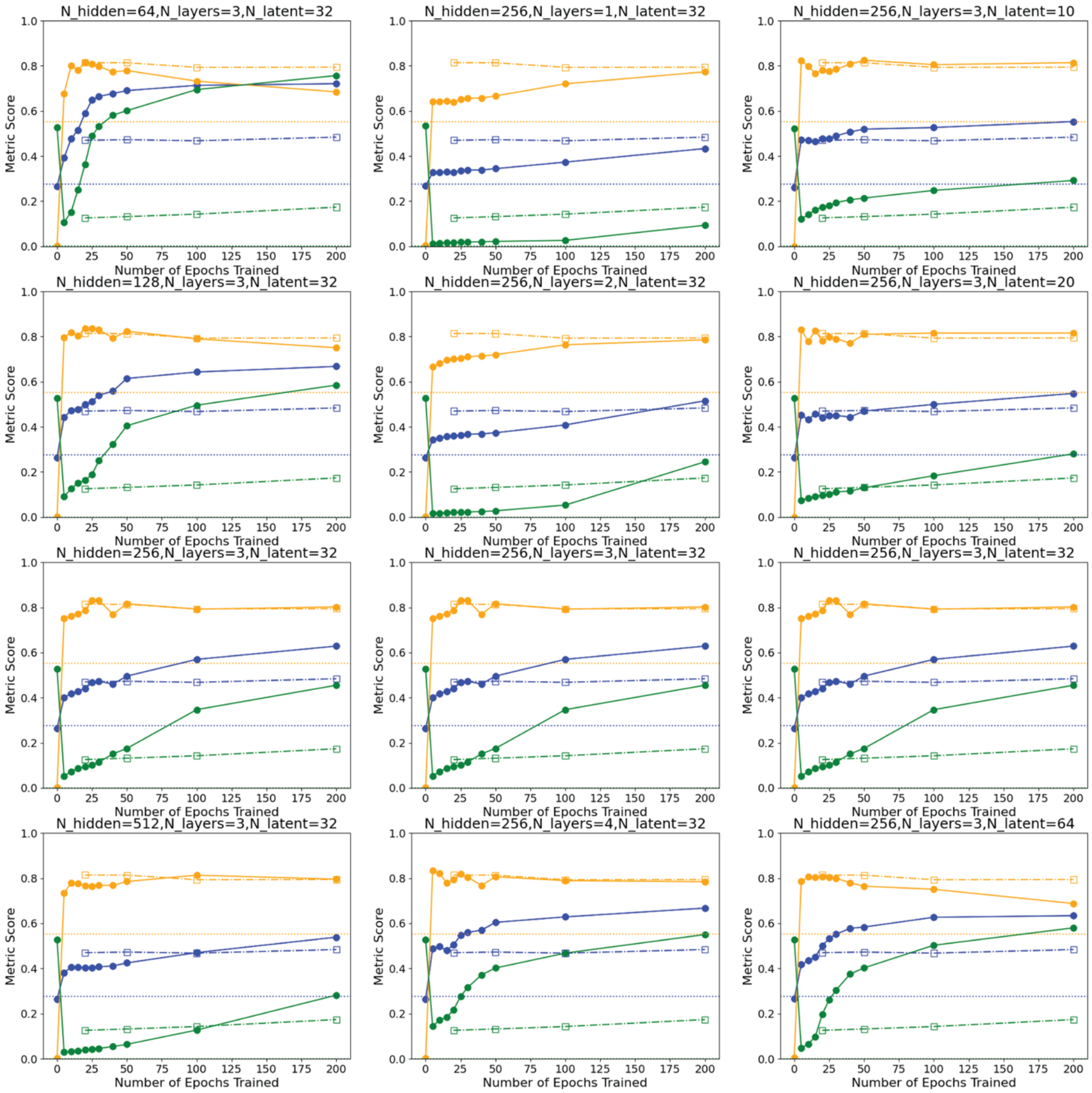


Figure S6: Benchmarking model architectures on primate basal ganglia integration. PIANO performs well with a default model architecture with 3 hidden layers, 256 hidden nodes per hidden layer, and 32 latent dimensions compared to other model architectures. scVI with 3 hidden layers, 256 hidden nodes per hidden layer, and 32 latent dimensions at 200 epochs is shown as a comparison.

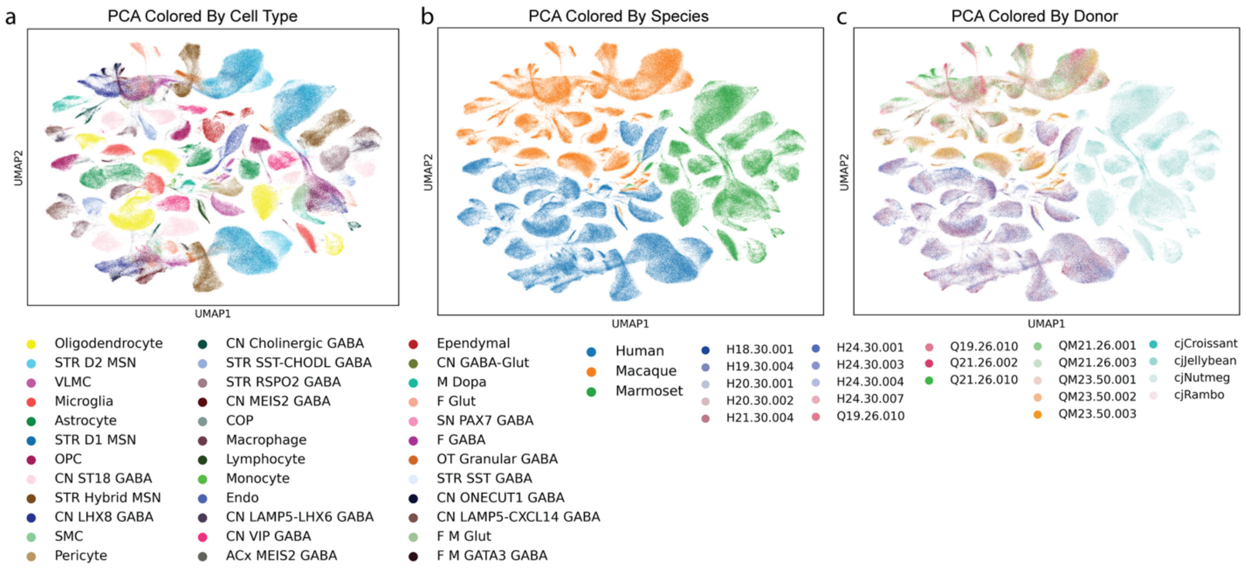


Figure S7: Principal component analysis of primate basal ganglia. Uniform manifold approximation and projections (UMAPs) of principal component analysis (PCA) of data colored by Subclass (a), species (b), and donor (c).

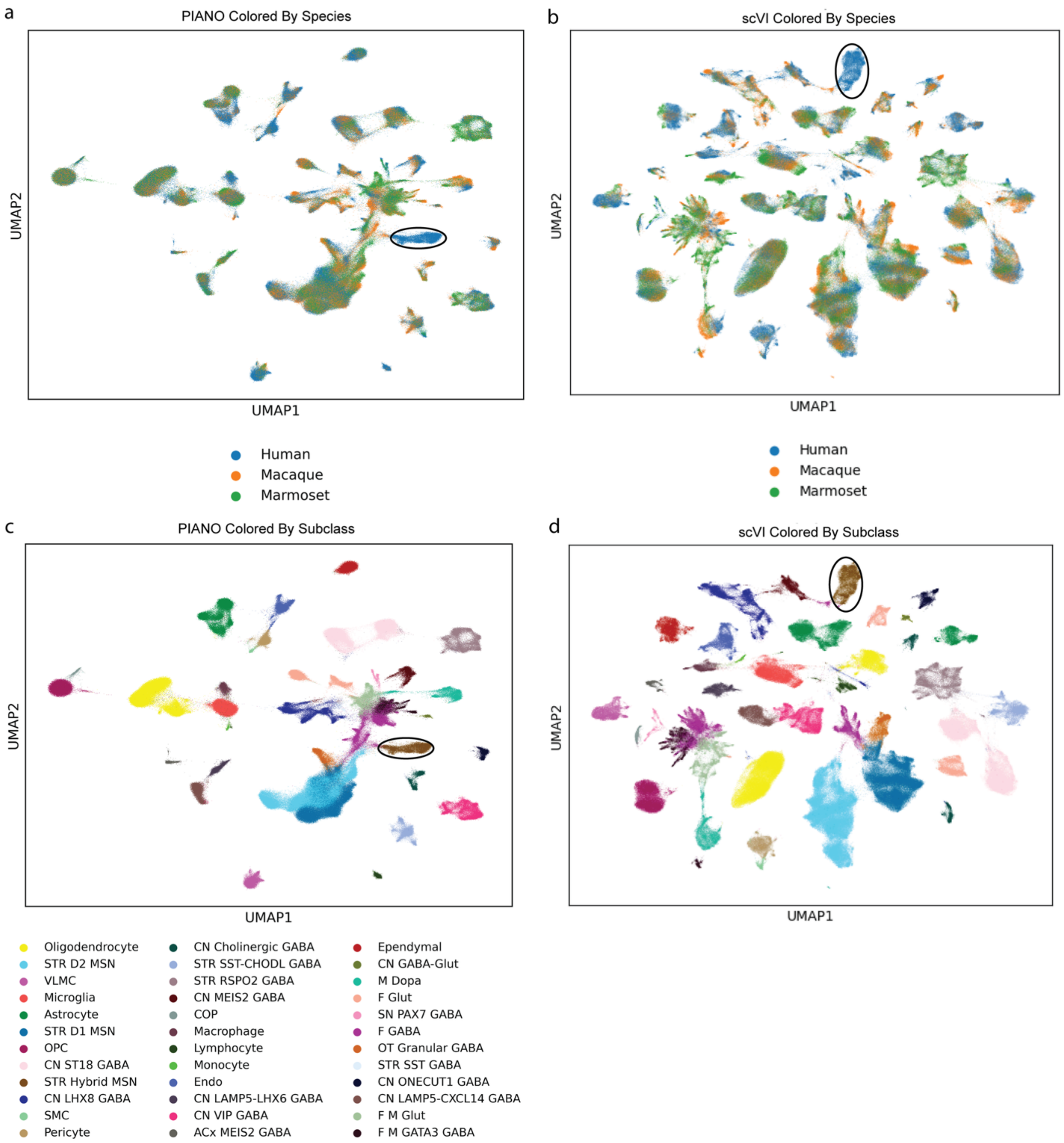


Figure S8: Primate basal ganglia integration with perturbation of withholding non-human primate Hybrid MSNs. PIANO and scVI preserve the perturbed human Hybrid MSNs as seen in uniform manifold approximation and projections (UMAPs) with the Hybrid MSN circled and colored by species for PIANO (a) and scVI (b) and colored by Subclass for PIANO (c) and scVI (d).

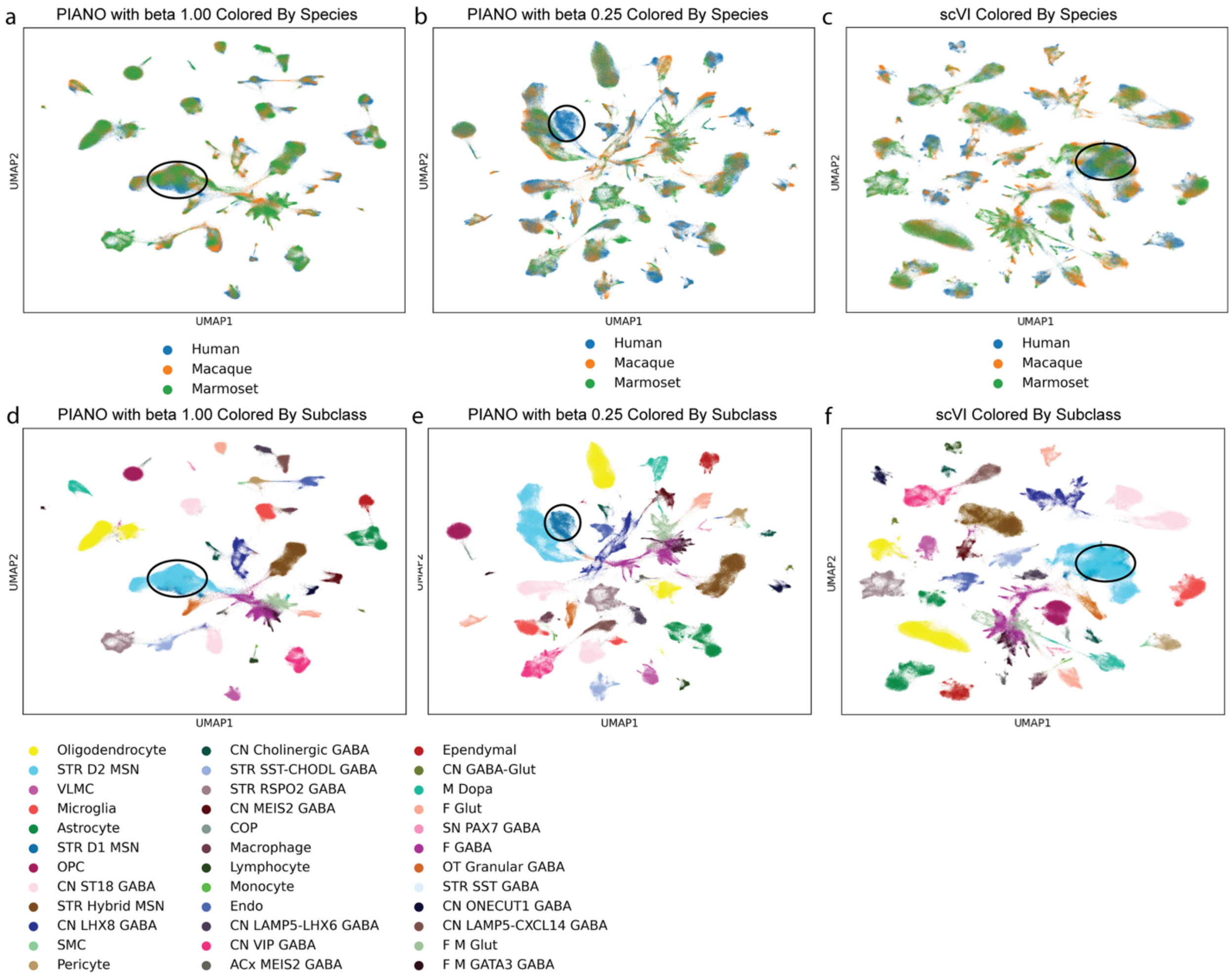


Figure S9: Primate basal ganglia integration with perturbation of withholding non-human primate D1 MSNs. PIANO with default parameters mixes the perturbed human D1 MSNs (a) but not with the beta parameter set to 0.25 (b). scVI mixes the perturbed human D1 MSNs (c). The respective uniform manifold approximation and projections (UMAPs) for PIANO and scVI are colored by Subclass (d-f).

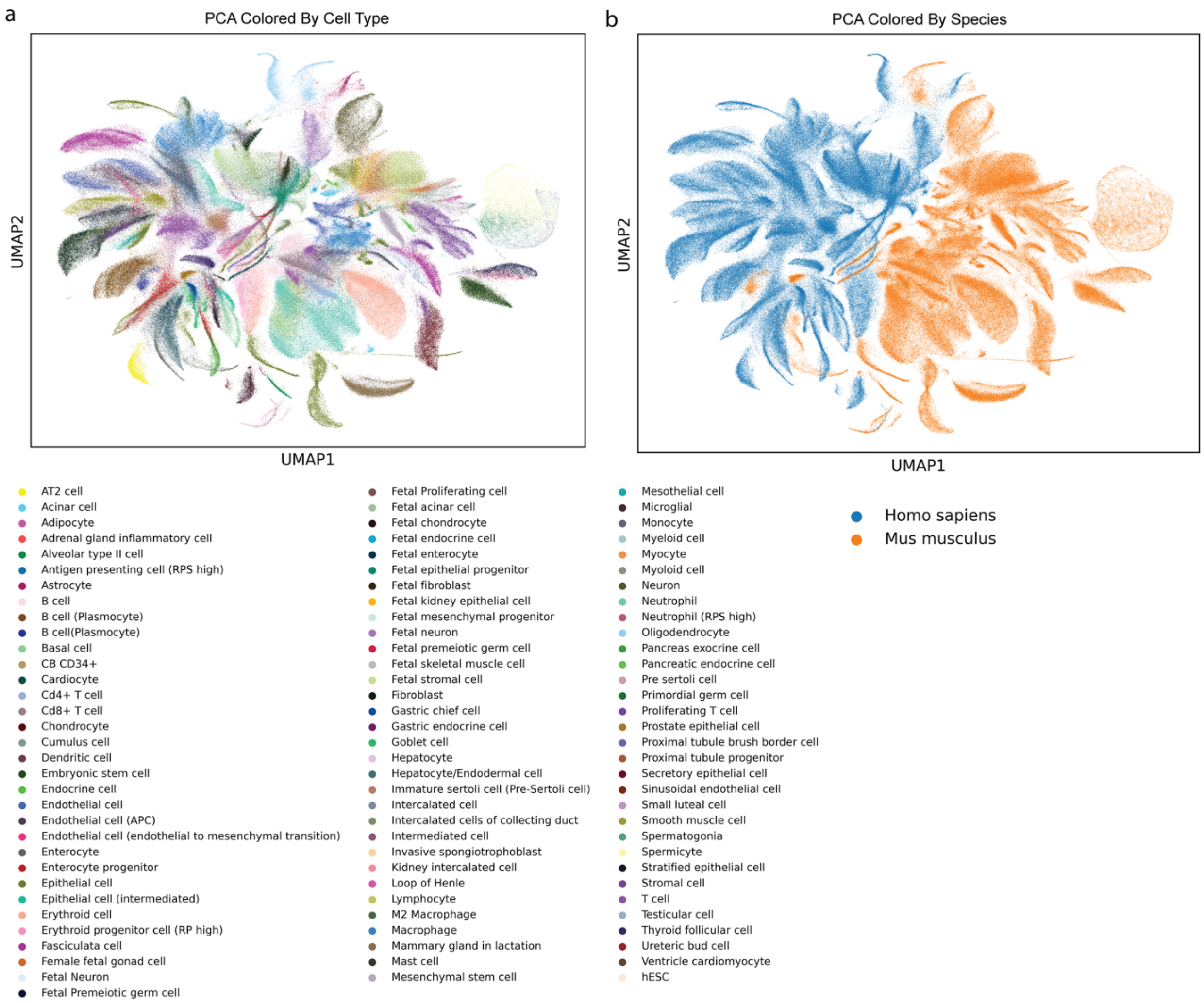


Figure S10: Principal component analysis of mouse cell atlas and human cell landscape. Uniform manifold approximation and projections (UMAPs) of principal component analysis (PCA) of mouse cell atlas and human cell landscape colored by cell type (a) and species (b).

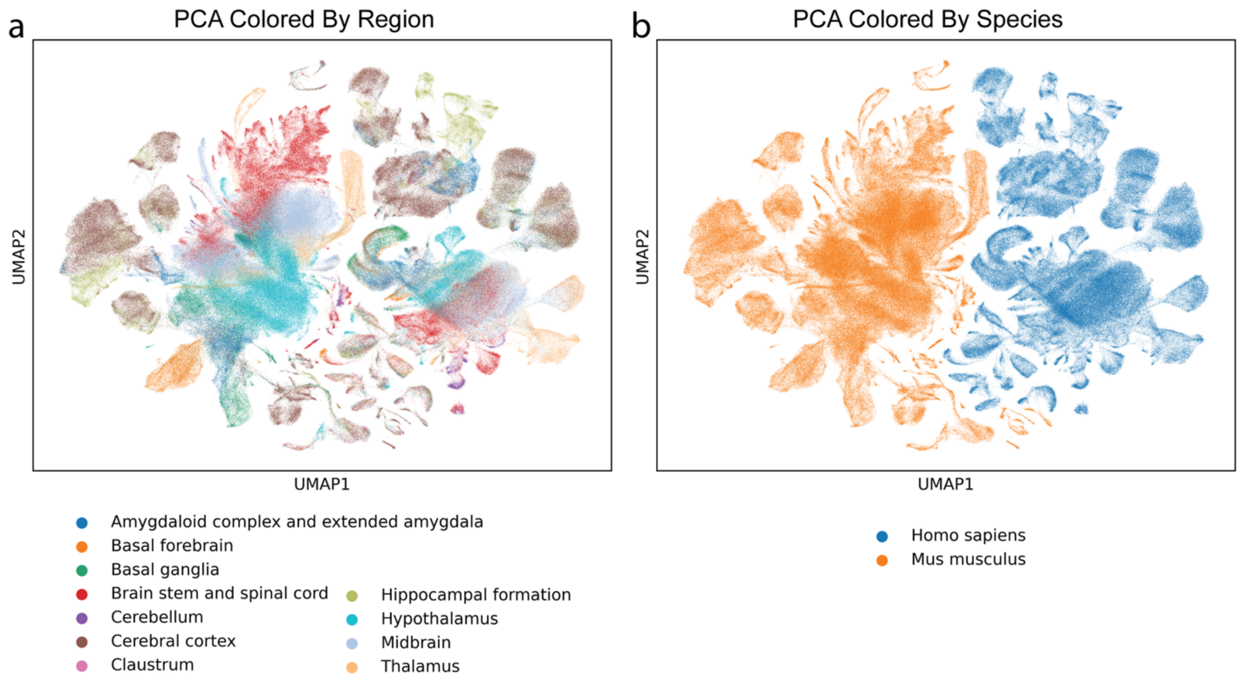


Figure S11: Principal component analysis of whole mouse and human brain atlases. Uniform manifold approximation and projections (UMAPs) of principal component analysis (PCA) of mouse and human whole brain atlases colored by region (a) and species (b).

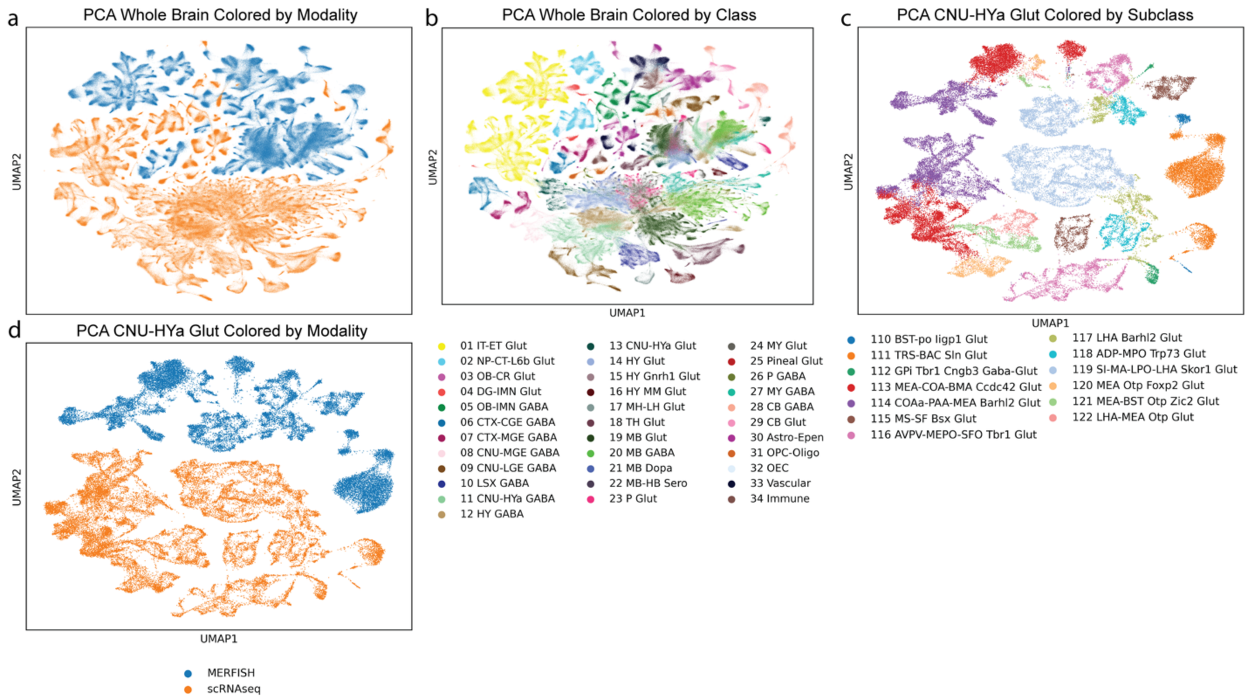


Figure S12: Principal component analysis of whole mouse brain single cell and spatial atlases. Uniform manifold approximation and projections (UMAPs) of principal component analysis (PCA) of whole mouse brain 10X scRNAseq and MERFISH spatial transcriptomics data colored by modality (a) and Class (b) and Class 13 CNU-HYa Glut colored by modality (c) and Subclass (d).

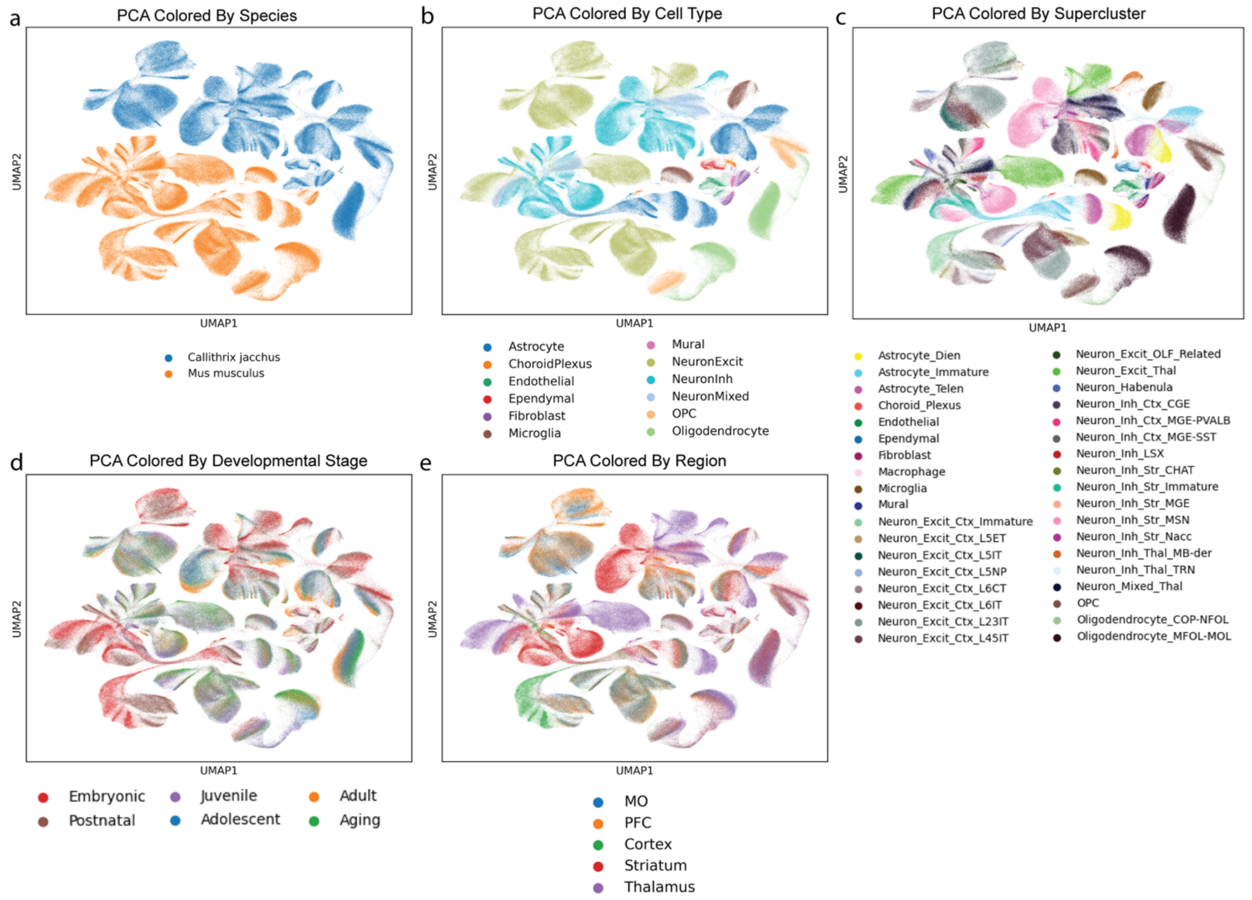


Figure S13: Principal component analysis of mouse and marmoset brain developmental data. Uniform manifold approximation and projections (UMAPs) for principal component analysis (PCA) of mouse and marmoset developmental data colored by species (a), cell type (b), supercluster (c), developmental time point (d), and brain region (e).
