## Supplementary material for "Fast Optimization of Robust Transcriptomics Embeddings using Probabilistic Inference Autoencoder Networks for multi-Omics": Key Resources Table

| REAGENT or RESOURCE | SOURCE | IDENTIFIER |
| --- | --- | --- |
| Deposited data | | |
| Lung data | Vieira Braga et al.^11^ | <https://figshare.com/articles/dataset/Benchmarking_atlas-level_data_integration_in_single-cell_genomics_-_integration_task_datasets_Immune_and_pancreas_/12420968?file=24539828> |
| Pancreas data | Grün et al.^12^ Muraro et al.^13^ Lawlor et al.^14^ Segerstolpe et al.^15^ | <https://figshare.com/articles/dataset/Benchmarking_atlas-level_data_integration_in_single-cell_genomics_-_integration_task_datasets_Immune_and_pancreas_/12420968?file=24539828> |
| Heart data | Kanemaru et al.^16^ | <https://figshare.com/articles/dataset/Batch_Alignment_of_single-cell_transcriptomics_data_using_Deep_Metric_Learning/20499630/2> |
| Immune data | Oetjen et al.^17^ Datasets - Single Cell Gene Expression - Official 10x Genomics Support^18^ Freytag et al.^19^ Sun et al.^20^ Villani et al.^21^ | <https://figshare.com/articles/dataset/Benchmarking_atlas-level_data_integration_in_single-cell_genomics_-_integration_task_datasets_Immune_and_pancreas_/12420968?file=24539828> |
| Tabula muris data | Tabula Muris Consortium^22^ | <https://figshare.com/articles/dataset/Single-cell_RNA-seq_data_from_Smart-seq2_sequencing_of_FACS_sorted_cells_v2_/5829687/8> |
| Primate Basal Ganglia data | Human and Mammalian Brain Atlas Consortium | [https://alleninstitute.github.io/HMBA_BasalGanglia_Consensus_Taxonomy/#rna-seq-data](https://alleninstitute.github.io/HMBA_BasalGanglia_Consensus_Taxonomy/" \l "rna-seq-data) |
| Ortholog gene tables | Dyer et al.^23^ | https://mart.ensembl.org/index.html |
| UniProtKB gene lists | Ahmad et al.^24^ | https://www.uniprot.org/uniprotkb |
| AnimalTFDB4 gene lists | Shen et al.^15^ | https://guolab.wchscu.cn/AnimalTFDB4/#/Download |
| Mouse Cell Atlas data | Han et al.^28^ | <https://figshare.com/articles/dataset/HCL_DGE_Data/7235471> |
| Human Cell Landscape data | Han et al.^29^ | <https://figshare.com/articles/dataset/HCL_DGE_Data/7235471> |
| Allen Institute whole mouse brain atlas single cell RNA sequencing data | Yao et al.^2^ | <https://alleninstitute.github.io/abc_atlas_access/descriptions/WMB_dataset.html> |
| Broad Institute whole mouse brain MERFISH data | Zhang et al.^31^ | <https://alleninstitute.github.io/abc_atlas_access/descriptions/Zhuang_dataset.html> |
| Karolinska Institute whole human brain data | Siletti et al.^1^ | <https://cellxgene.cziscience.com/collections/283d65eb-dd53-496d-adb7-7570c7caa443> |
| Developmental mouse data | Schroeder et al.^27^ | <https://singlecell.broadinstitute.org/single_cell/study/SCP2719/a-multi-region-transcriptomic-atlas-of-developmental-cell-type-diversity-in-mouse-brain> |
| Developmental marmoset data | Schroeder et al.^27^ | <https://singlecell.broadinstitute.org/single_cell/study/SCP2706/a-multi-region-transcriptomic-atlas-of-developmental-cell-type-diversity-in-marmoset-brain> |
| Software and algorithms | | |
| Seurat 5.3.0 (Using Seurat V3 CCA) | Butler et al.^6^ Stuart et al.^32^ | <https://satijalab.org/seurat/> |
| Harmony 0.0.10 | Korsunsky et al.^5^ | <https://portals.broadinstitute.org/harmony/> |
| scIB benchmarking metrics | Luecken et al.^7^ | [https://scib-metrics.readthedocs.io/en/stable/#installation](https://scib-metrics.readthedocs.io/en/stable/" \l "installation) |
| scvi-tools v1.3.3 | Lopez et al.^9^ | <https://docs.scvi-tools.org/en/stable/installation.html> |
| ScanPy 1.11.4 | Wolf et al.^40^ | <https://scanpy.readthedocs.io/en/stable/installation.html> |
